# Multiscale 3D whole joint cellular and molecular mapping dissects the relationship between structure and pain

**DOI:** 10.1101/2025.08.23.671843

**Authors:** Peng Chen, Jiaxin Chai, Abirami Soundararajan, R. Glenn Hepfer, Benjamin Kheyfets, Jiaxin Hu, Ishraq Alshanqiti, Swarnalakshmi Raman, Ikue Tosa, Jun Tae Huh, Matthew Yee, Brooke J. Damon, Shangping Wang, Yu Shin Kim, Man-Kyo Chung, Mildred C. Embree, Janice S. Lee, Tong Ye, Hai Yao

## Abstract

Understanding musculoskeletal joints from a 3D multiscale perspective, from molecular to anatomical levels, is essential for resolving the confounding relationships between structure and pain, elucidating the intricate mechanisms regulating joint health and diseases, and developing new treatment strategies. Here, we introduce a musculoskeletal joint immunostaining and clearing (MUSIC) method specifically designed to overcome key challenges of immunostaining and optical clearing of intact joints. Coupled with large-field light sheet microscopy, our approach enables 3D high-resolution, microscale neurovascular mapping within the context of whole-joint anatomy without the need for image coregistration across various joints, including temporomandibular joints, knees, and spines, and multiple species, including mouse, rat, and pig. Our findings reveal 3D heterogeneous distributions of neurovascular networks and previously uncharacterized neurovascular pathways within joints. Using the proteoglycan 4 knockout (*Prg4^-/-^*) mouse model of joint degeneration, we identified significant alterations in joint-wide neurovascular architecture, highlighting neurovascular changes along degenerative processes. Furthermore, in a traumatic joint injury mouse model, we observed long-lasting pain behavior and a time-course 3D neurovascular remodeling preceding detectable joint morphological change, bridging microscale alterations with potential pain mechanisms. This platform offers a powerful tool for multiscale 3D analysis, enabling new insights into joint pathophysiology and intricate interplay among joint tissues.

## INTRODUCTION

Musculoskeletal diseases, such as temporomandibular joint disorders, osteoarthritis, and back pain, are the leading cause of disability, affecting approximately 1.7 billion people worldwide and resulting in an estimated annual healthcare cost of $980 billion in the United States alone ^1,2^. The musculoskeletal joint is a diverse and multiscale system comprising an array of different joint tissues (i.e., bones, connective tissues, and muscles), each with unique 3D structural and functional characteristics that span from the anatomical to molecular levels ^3,4^. Historically, different approaches have often been utilized to assess the joint systems at a given scale. For instance, magnetic resonance imaging (MRI) is employed to evaluate macroscale soft tissue anatomy and composition, computed tomography (CT) is used to visualize bony structures, and histological techniques provide microscale insights into cellular and molecular markers. However, interpreting joint diseases through a single imaging modality can be confounding. One prominent example is the long-standing challenge to dissect the relationship between joint structure alterations, dysfunction, and pain ^5,6^. Radiographic images of macroscale joint abnormalities do not consistently align with reported pain levels, complicating diagnosis and impeding the development of effective, comprehensive treatments ^5,6^. This discordance underscores the need for an integrated, multiscale approach to assessment.

Recent advancements in imaging have explored multiscale visualization of 3D joint vasculature by integrating MRI, CT, and optical imaging ^7^. While promising, this approach necessitates multiple scans and complex image coregistration across modalities, and it remains primarily limited to vascular mapping due to its dependence on vessel-based contrast agents ^7^. Although the vascular system is vital for maintaining joint integrity and homeostasis by facilitating oxygen and nutrient delivery, molecular signaling, and cellular trafficking ^8,9^, understanding the relationship between joint structure and pain necessitates the 3D mapping of additional cellular and molecular components, particularly the neural network ^10,11^. Nerve fibers innervate the joints to sense the intrinsic and environmental stimuli (e.g., proprioceptive and noxious stimuli), transmit the signals to the brain where pain is perceived, and trigger somatic responses ^11^. Beyond their sensory role, neural elements also contribute to joint development and remodeling through the release of a variety of neurotrophic and regulatory factors ^12^. Therefore, there is a critical need for next-generation 3D imaging techniques with improved molecular specificity that can capture the multiscale interplay among vascular, neural, and other tissue-specific elements.

Optical imaging methods, especially fluorescence-based ones, allow high-resolution characterization of various cell and tissue structures owing to the availability of many molecular-specific endogenous and exogenous fluorescent probes. However, 3D imaging large or thick samples presents a significant challenge due to the limited light penetration caused by tissue scattering, especially along the optical axial dimension. While multiphoton imaging can extend imaging depth to hundreds of micrometers ^13^, it remains insufficient for multiscale 3D whole-joint imaging. To overcome these limitations, two primary strategies have been employed: serial section reconstruction and tissue clearing. In the serial section reconstruction, the sample is physically cut into a series of thin sections and imaged section by section for 3D reconstruction; this method is labor-intensive, technique sophisticated, and subject to sample distortion due to the invasive sample sectioning ^14–17^. Tissue clearing techniques offer a powerful alternative by rendering tissues optically transparent, thereby preserving their native 3D architecture and allowing for deeper and multiscale imaging ^18,19^. Over the past decade, many tissue clearing protocols and methods have been developed ^19^, with the majority applied in the brain and increasing applications in other organs, including musculoskeletal tissues ^20–26^. However, previous reports only examined single joint components in mice, such as muscles ^21,27^ or bones ^20,22–25,28,29^. Systematic whole joint mapping presents significant challenges due to the complex architecture of musculoskeletal joints ^30^. Firstly, unlike the brain which contains only soft tissues, musculoskeletal joints uniquely have hard and soft components, each exhibiting unique properties such as structure, composition, and optical transparency ^30^. Secondly, tissues in the joint, like the tendon and meniscus, are very dense with high collagen and glycosaminoglycan (GAG) contents, posing difficulties for antibody labeling and tissue clearing for the whole joint, as reported in recent publications ^30,31^. Thirdly, the diverse array of joint types, including knee joints, spines, and TMJs, necessitates a flexible approach accommodating different anatomical structures. Lastly, the size variability of joints in commonly utilized preclinical models, including mice, rats, and pigs, ranges from millimeters to centimeters, which underscores the need for scalable tissue clearing techniques and large-field 3D imaging methods. A platform method with broad utility for 3D multiscale whole joint mapping across various joint types, sizes, and species has yet to be developed.

Here, we present a **mus**culoskeletal joint **i**mmunostaining and **c**learing technique (**MUSIC**), which specifically addresses key challenges of immunostaining and optical clearing of intact joints with various types and sizes through decalcification, deep joint permeabilization with small-micelle detergent and GAGs extraction, and antibody stabilization. Coupled with fast and large-field light sheet microscopy, our approach enables 3D whole-joint cellular and molecular mapping in a range of joints—including TMJs, knees, and spines—across multiple species, including mice, rats, and pigs. Our method allows multiscale 3D joint imaging of both macroscale joint anatomy and microscale cellular and molecular features, while no image coregistration is needed. Using the mouse TMJ as an example, we first demonstrated successful immunostaining, clearing, and imaging of the 3D microscale neurovascular networks within the entire intact joint under healthy and diseased conditions. Our results illustrated the heterogeneous distribution of vessels and nerves in healthy TMJ and revealed detailed pain-sensing nerve fiber innervation pathways. Using a well-established mouse model of degenerative joint disease with global deficiency in proteoglycan 4 (*Prg4^-/-^*) ^32,33^, we revealed iconic TMJ structure degeneration from a 3D perspective that was coupled with a drastically vascularized and innervated lateral capsule and synovial membrane and altered condyle vasculature networks that have not been shown previously. We further uncovered the time-course 3D plasticity of neurovascular structure in a TMJ injury mouse model, the forced-mouth opening (FMO) model ^34^. We observed distinct microscale temporal degenerative changes of pain-sensing nerves within the injured anterior region of the TMJ disc, despite the absence of detectable macroscale alterations in joint morphology or anatomy. Aligned with pain behavior assessments, our data provides valuable new insights to bridge the joint structure changes with pain. Additionally, we have demonstrated our method in mouse knees and found sprouting of 3D neurovascular structures in the knee capsules of *Prg4^-/-^* mice compared to wild-type mice. Our method was further scaled up to map the 3D neurovascular structures in the whole rat TMJ, knee, and spine samples and pig TMJ discs with surrounding tissue attached. Overall, we provided a scalable and versatile tissue clearing and imaging method for multiscale 3D whole joint cellular and molecular mapping in musculoskeletal systems. This method allows us to quantitatively dissect the relationship between joint structure and pain alongside disease progression, repair, and following therapy, thereby advancing mechanistic studies and treatment development.

## RESULTS

### MUSIC for whole joint cellular and molecular mapping

Tissue clearing involves a series of chemical treatments to homogenize tissues ^19^. Selecting tissue clearing protocols highly depends on how the targeted molecules are labeled. For instance, endogenous fluorescent proteins need a delicate strategy to preserve their fluorescence ^25,30,35,36^. In general, there are two approaches for molecular labeling: endogenous fluorescent proteins based on transgenic animals or exogenous fluorescent probes through immunostaining or chemical labeling. Compared to the transgenic approach, immunostaining or chemical labeling has the advantages and flexibility to simultaneously study multiple cellular or molecular targets across different species at a reasonable price, benefiting from the large pool of antibodies and probes on the market ^35,37,38^. Considering our intention to develop a versatile platform method that works for different research targets, such as investigating the neural and vascular structure and subtypes of the nerves and vessels, we decided to focus on the immunostaining method for target molecular labeling.

To achieve whole joint immunostaining and clearing, a new pipeline was developed based on the unique joint features. First, the fixed joints were decalcified and then treated with a deep permeabilization chemical, CHAPS (3-[(3-cholamidopropyl) dimethylammonio]-1- propanesulfonate). CHAPS is a zwitterionic detergent that can form much smaller micelles than traditional tissue clearing detergents, like Triton X-100, allowing rapid and efficient tissue permeabilization, as recently demonstrated with human brain tissues ^31^. Next, guanidine hydrochloride was used to remove the glycoproteins in joint tissues with high GAGs while keeping the fixed collagen backbones to achieve extracellular matrix loosening ^31^, enhancing the immunostaining antibody penetration. Given the various sizes of joints across different species, ranging from 5 mm to 5 cm, several days of antibody incubation are still needed, even with our efforts to deeply permeabilize the samples. Antibodies aggregate during the prolonged staining process, causing problems for labeling, visualization, and data analysis ^35^. To mitigate this problem, we used heptakis(2,6-di-O-methyl)-β-cyclodextrin, a chemical that has been recently proven to reduce antibody aggregation ^35^. Finally, the organic solvent, dibenzyl ether (DBE), was selected as a refractive index matching agent for tissue clearing, given its superior compatibility with immunostaining and its excellent performance in clearing complicated organs, including musculoskeletal bones and muscles ^28,36–38^. With optimizations based on the unique features of musculoskeletal joints, we finalized the MUSIC protocol for the whole joint mapping (**Fig. 1a**). To achieve fast and large-field imaging, an advanced light sheet microscope ^39^ was applied for whole joint mapping (**Fig. 1a**). Compared to conventional confocal microscope imaging that detects fluorescence signal in a pixel-based manner with a low quantum efficient photomultiplier tube, light sheet microscope uses a sheet of light illuminating a layer of sample and captures information from the entire layer at once using a highly sensitive camera, significantly increasing the 3D imaging speed and reducing photobleaching effects ^40^. The light sheet microscope used in this study is also equipped with a special lens that provides a large field of view and a dynamic scanning algorithm optimized for uniform illumination of large samples. The sample volume can be imaged up to a volume of 53 cm^3^, which covers the dimensions of a wide range of joint samples.

**Figure 1.**
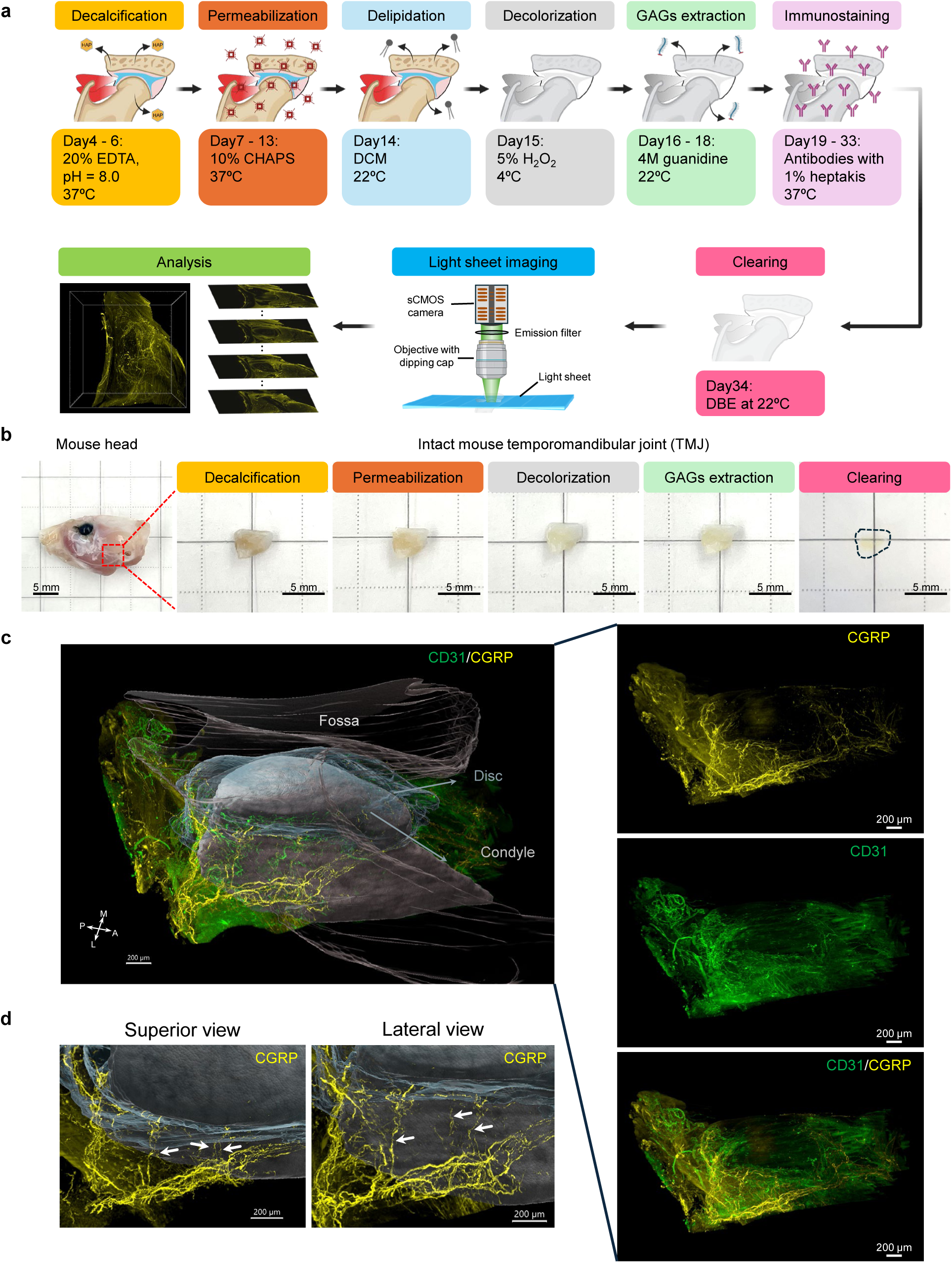
MUSIC for whole joint neurovascular mapping in mouse TMJ. **a.** Schematic workflow and timeline for the MUSIC method with mouse TMJ samples. Fixed joint samples were first decalcified in 20% EDTA solution and then incubated in 10% CHAPS solution for deep permeabilization. Then, they are dehydrated, delipidated with DCM, and decolored with 5% H_2_O_2_ solution. Next, the joint samples were treated with 4M guanidine solution to extract the GAGs from the joint tissue extracellular matrix while maintaining their collagen backbone. Then, the joints were stained with primary and secondary antibodies (**Supplementary Table 1**) immunostaining solution with the addition of 1% heptakis(2,6-di-O-methyl)-β-cyclodextrin to minimize antibody aggregation. Finally, the joints were cleared in DBE and imaged with a large-field light sheet microscope. The 3D mapping data were further reconstructed and analyzed. Similar procedures were applied to other joints (**Supplementary Table 2**). **b.** Photos of mouse head and mouse TMJs along MUSIC procedures. The red dashed square highlighted the region where the TMJ was harvested. The black dashed line depicted the contour of a cleared mouse TMJ. Scale bar, 5 mm. **c.** Whole joint neurovascular mapping in a mouse TMJ. The TMJ condyle, disc, and fossa were segmented, reconstructed, and rendered in grey, translucent light blue, and translucent light brown, respectively, to provide a joint coordinate to visualize the neurovascular structure distribution (**Supplementary Fig. 3a-3b**). 3D data from each imaging channel and merged channel without rendering were displayed on the right side. Signals from muscles were removed for visualization. Scale bar, 200 µm. **d.** Zoom-in view at the lateral condyle neck region of the mouse TMJ from superior and lateral angles. White arrows highlighted the nerve strands branching out from the major nerve bundles and innervating the TMJ disc. Scale bar, 200 µm. A, anterior; P, posterior; M, medial; L, lateral.

Using mouse TMJ as an example, we demonstrated successful whole-joint clearing using our method (**Fig. 1b**). With the staining of validated CD31 and calcitonin gene-related peptide (CGRP) antibodies for blood vessels and pain-sensing nerve fibers, respectively (**Supplementary Fig. 1**, **Supplementary Table 1**, and **Supplementary Note 1**), we achieved neurovascular mapping of the entire intact mouse TMJ (**Fig. 1c**). We further validated efficient whole joint immunostaining in mouse TMJs with the MUSIC method (**Supplementary Note 2** and **Supplementary Fig. 2**). Our method inherently captures the macroscale joint anatomy, allowing easy spatial registration of individual joint components (**Supplementary Fig. 3**). Through image segmentation, we can resolve the microscale neurovascular distribution in different joint tissues in the spatial context of the whole joint without the need of complicated image coregistration (**Fig. 1c**, **Supplementary Fig. 3a-3b**, and **Supplementary Movie 1**). Previous literature based on 2D histological analysis has reported innervation in the TMJ disc with inconsistent conclusions ^41^. Our results confirmed that no blood vessels and nerves are present in the central region of the disc (**Fig. 1c**). High density of neurovascular structures was observed in the anterior and posterior regions of the disc. In the zoomed-in view of the lateral side of the TMJ, strands of nerve fibers that branch from the major nerve bundles and innervate the disc were observed (**Fig. 1d**), unveiling the microscopic level of the innervation pathway of the TMJ disc. With our method, we can access the neurovascular spatial pattern in the entire TMJ that has never been seen before. It provides an unprecedented opportunity to explore its spatial distributions, tissue-specific features, and connections/interactions between each joint component.

### Whole joint degenerative changes in *Prg4^-/-^* mouse TMJ

Degenerative joint disease, such as osteoarthritis, is a major cause of musculoskeletal dysfunction and pain ^42^. Few studies have investigated the TMJ neurovascular structure in osteoarthritic joints, with inconsistent literature reports using 2D histological methods ^43^. Here, we applied our MUSIC method to investigate how the 3D neurovascular structure changes when joints degenerate. We used a well-established genetic mouse model globally deficient in *Prg4* ^32,33^. The *Prg4* gene encodes for the protein, lubricin, a critical glycoprotein in synovial joint fluid and cartilage superficial zone that provides joint lubrication ^32^. Mice lacking lubricin develop age-dependent osteoarthritis-like degeneration in joints, such as TMJs and knees ^32,33,44^. We first confirmed the joint degeneration in *Prg4^-/-^* mice through traditional hematoxylin and eosin (H&E) staining. The *Prg4^-/-^*TMJ showed a thickened disc, synovial membrane hyperplasia, articular cartilage erosion and fibrillation, and large marrow cavities, which matched well with literature observations ^32,33,44^ (**Fig. 2a**). TMJ bone morphometric analysis performed with micro-computed tomography (µCT) imaging showed that the *Prg4^-/-^* TMJ had calcified disc, deformed condyle, and condyle bone resorption (**Fig. 2b**). The *Prg4^-/-^* TMJ condyle 3D shape was rounder with significantly decreased length and increased width compared to wild-type mice (**Fig. 2b** and **Fig. 2c**). We confirmed similar pathological changes using our MUSIC mapping method relative to traditional histologic analysis in the *Prg4^-/-^* TMJ from a single 2D optical section (**Fig. 2d**). In addition, through 3D reconstruction and segmentation, we observed 3D condyle anatomical changes in the *Prg4^-/-^* TMJ as shown in the µCT analysis (**Fig. 2e**). More importantly, unlike H&E and µCT techniques, the MUSIC protocol allowed us to map the microscale neurovascular structure in the whole joint by immunostaining CGRP (a marker of peptidergic sensory nerves) and CD31 (a marker of blood vessels) and uncover a 3D highly vascularized and innervated lateral capsule and synovial membrane in the *Prg4^-/-^*TMJ (**Fig. 2e-2f**, **Supplementary Fig. 4a-4c**, and **Supplementary Movie 2-3**). The internal vasculature network can also be visualized with the integration of immunofluorescence and blood autofluorescence using the blood retention sample preparation approach ^45^ (**Fig. 2g**). The vasculature architecture dramatically changed within the *Prg4^-/-^* TMJ condyle head subchondral bone (**Fig. 2g** and **Supplementary Movie 4-5**), which may indicate aberrant subchondral bone remodeling. The pathways connecting the vasculature between the inner and outer regions of the condyle were also observed at the condyle neck (**Supplementary Fig. 5**), which could provide new insights into the crosstalk among different joint components.

**Figure 2.**
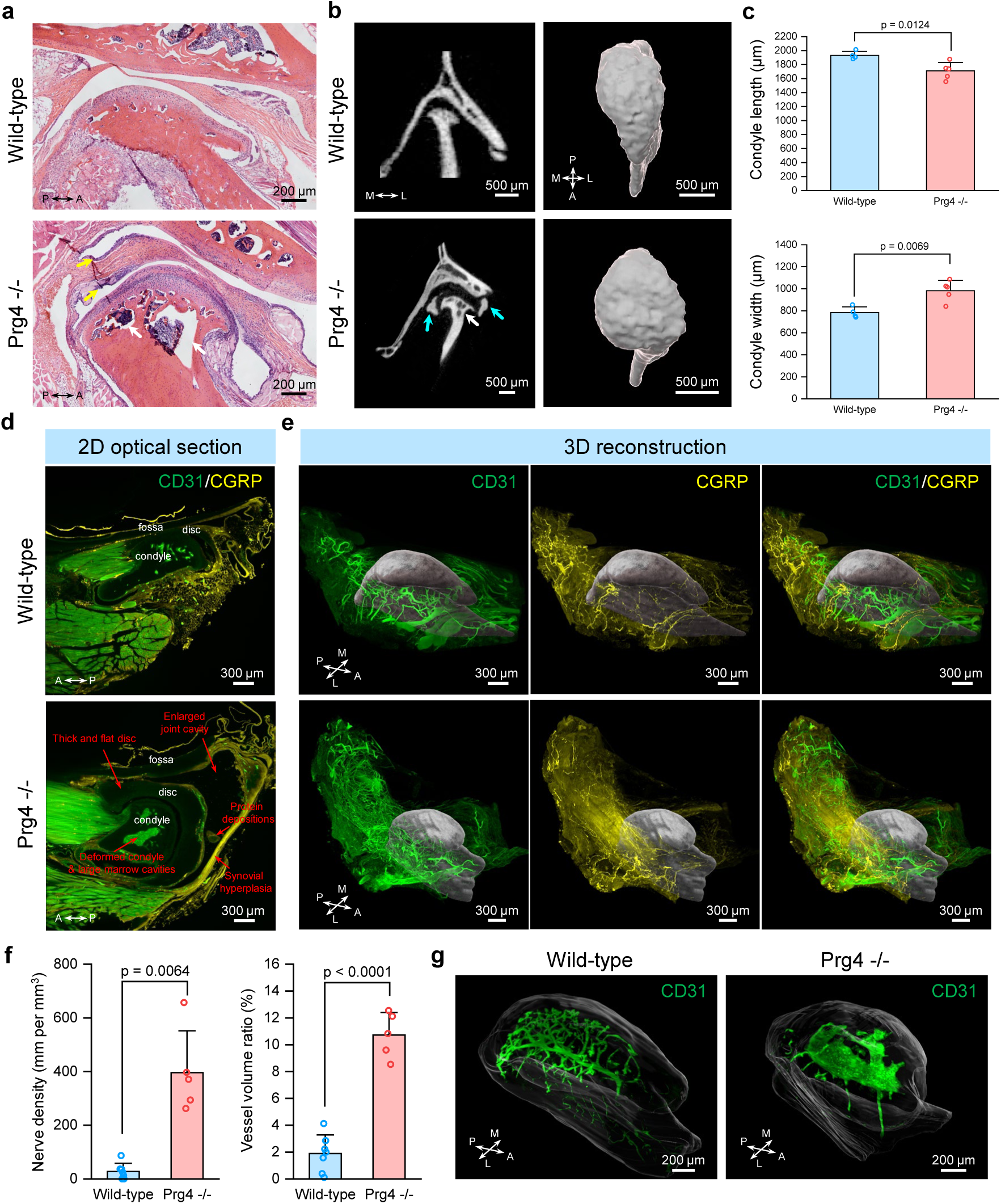
Whole joint neurovascular mapping in degenerative *Prg4^-/-^* mouse model. **a.** H&E staining results with mouse TMJs in wild-type and *Prg4^-/-^* mice. White arrows highlighted the large marrow cavities in the TMJ condyle head, and yellow arrows pinpointed the inflamed synovial membrane in *Prg4^-/-^* mice. Scale bar, 200 µm. **b.** µCT imaging results with mouse TMJs in wild-type and *Prg4^-/-^* mice. Left, 2D µCT images of mouse TMJ. White arrows highlighted the large marrow cavities in the TMJ condyle head, while the cyan arrows depicted the TMJ disc calcification in *Prg4^-/-^* mice. Right, 3D reconstruction of mouse TMJ condyle head. Scale bar, 500 µm. **c.** Quantitative TMJ condyle head size comparison between wild-type (n = 4 measurements from 4 mice) and *Prg4^-/-^* mice (n = 5 measurements from 5 mice). p-value was determined with a two-sided t-test. All data depict mean ± standard deviation. **d.** 2D optical section from the 3D neurovascular mapping in the whole mouse TMJ in wild-type and *Prg4^-/-^* mice. TMJ condyle, disc, and fossa are labeled. The red arrows highlighted the pathological changes in the TMJ of *Prg4^-/-^*mice. Scale bar, 300 µm. **e.** 3D reconstruction results of whole joint neurovascular mapping in mouse TMJ of wild-type and *Prg4^-/-^* mice. The TMJ condyles were segmented from the mapping data and rendered in grey to provide a spatial reference. Scale bar, 300 µm. **f.** Quantitative nerve and blood vessel density in wild-type (n = 7 measurements from 4 mice) and *Prg4^-/-^* TMJs (n = 5 measurements from 3 mice). The nerve and vessel densities were quantified at the lateral capsule with synovial membrane (**Supplementary Fig. 4a-4c**). p-value was determined by a two-sided t- test. All data depict mean ± standard deviation. **g.** 3D reconstruction of vascular structures within the TMJ condyle head. The TMJ condyle was segmented and rendered in translucent color to provide a spatial reference. Scale bar, 200 µm. A, anterior; P, posterior; M, medial; L, lateral.

In summary, using the *Prg4^-/-^* mouse model, we demonstrated the capacity of our method for multiscale whole joint neurovascular mapping and its application in joint disease research. The beauty of our method is that the entire 3D joint information is collected, including soft and hard tissue anatomy and cellular and molecular patterns. Existing imaging tools cannot fully recapitulate joint structures, particularly soft tissues, even with MRI, due to resolution limits. Using our method, we can obtain multimodality data from a single platform. Our approach provides a powerful tool to investigate multiple joint components simultaneously and unveil their crosstalk and contributions to joint health and diseases, supporting the whole joint research concept.

### Spatiotemporal neurovascular remodeling in FMO injury mouse model links structure and pain

TMJ is one of the most frequently used load-bearing joints. Overloading the joint can cause joint injury, pain, and dysfunction ^34,46^. TMJ injury animal models, such as the forced-mouth opening (FMO) model, have been developed to investigate the joint’s structural response to biomechanical stimulus and how it relates to pain and joint function ^34,46^. With our whole joint mapping approach, we can holistically study this question. Adult mice (8 – 12 weeks old) were subjected to either FMO or Sham procedures under anesthesia for 3 hours per day for five consecutive days, as described in previous publications ^34^ (**Fig. 3a**). Sham mice were sacrificed 12 days after the procedure, while FMO mice were euthanized at multiple time points post-injury (12 days (FMO_12d), 26 days (FMO_26d), and 58 days (FMO_58d)) (**Fig. 3a**). Our recently published pain behavior data has demonstrated that FMO induces long-last mechanical pain ^34^ (**Fig. 3b**). However, histologically, we did not see apparent differences among groups in the cartilage, synovial, or condyle bone (**Fig. 3c**). 3D µCT analysis of condyle size also showed no significant differences among the groups (**Fig. 3d** and **3e**). In contrast, time-course neurovascular structure changes in the TMJ were observed through our 3D mapping data (**Fig. 3f** and **Supplementary Movie 6-9**). We quantified the nerve and blood vessel density in the anterior region of the TMJ disc (**Supplementary Fig. 4d-4e**), where tissue injury happened during the FMO procedure ^47,48^. Our results showed that the nerve density was significantly reduced following the injury and then gradually increased over time, even though it did not return to the same level as the Sham group (**Fig. 3g**). Such neural changes indicate a dynamic process of nerve terminal degeneration and regeneration, similar to what has been reported in skin wound healing ^49^. These data may help explain our pain behavior results of sustained pain in FMO mice (**Fig. 3b**) ^34^. It is possible that the remaining nerve fibers in the TMJ discs get sensitized after injury, and/or the nerve terminal injury causes neuropathic-like persistent pain ^48,50^. Although the pain sources in TMJ remain elusive, a more detailed investigation of neural changes in the TMJ should provide new insights into peripheral pain-sensing mechanisms. Whole joint neural mapping provides the anatomic or structural basis for further investigation of joint pain. Interestingly, the blood vessel density showed no significant differences among the groups, with only a slight increase in the FMO_12d group (**Fig. 3h**). The disparity between the neural and vascular structure changes suggests that vascularity shows higher resistance to tissue injury or blood vessel ingrowth preceding innervation during wound healing. Our data uncovered significant neurovascular changes before any apparent changes in joint anatomy (**Fig. 3c-3e**), demonstrating the sensitivity of our method to detect microscopic levels of cellular and molecular alterations. In addition, it has been controversial to correlate joint structure changes (i.e., joint morphology through CT or MRI) and joint symptoms (e.g., pain) ^6^. Our results offer a new data dimension, providing microscopic-level information from a whole joint perspective to deepen our understanding of the joint’s structure-pain relationship.

**Figure 3.**
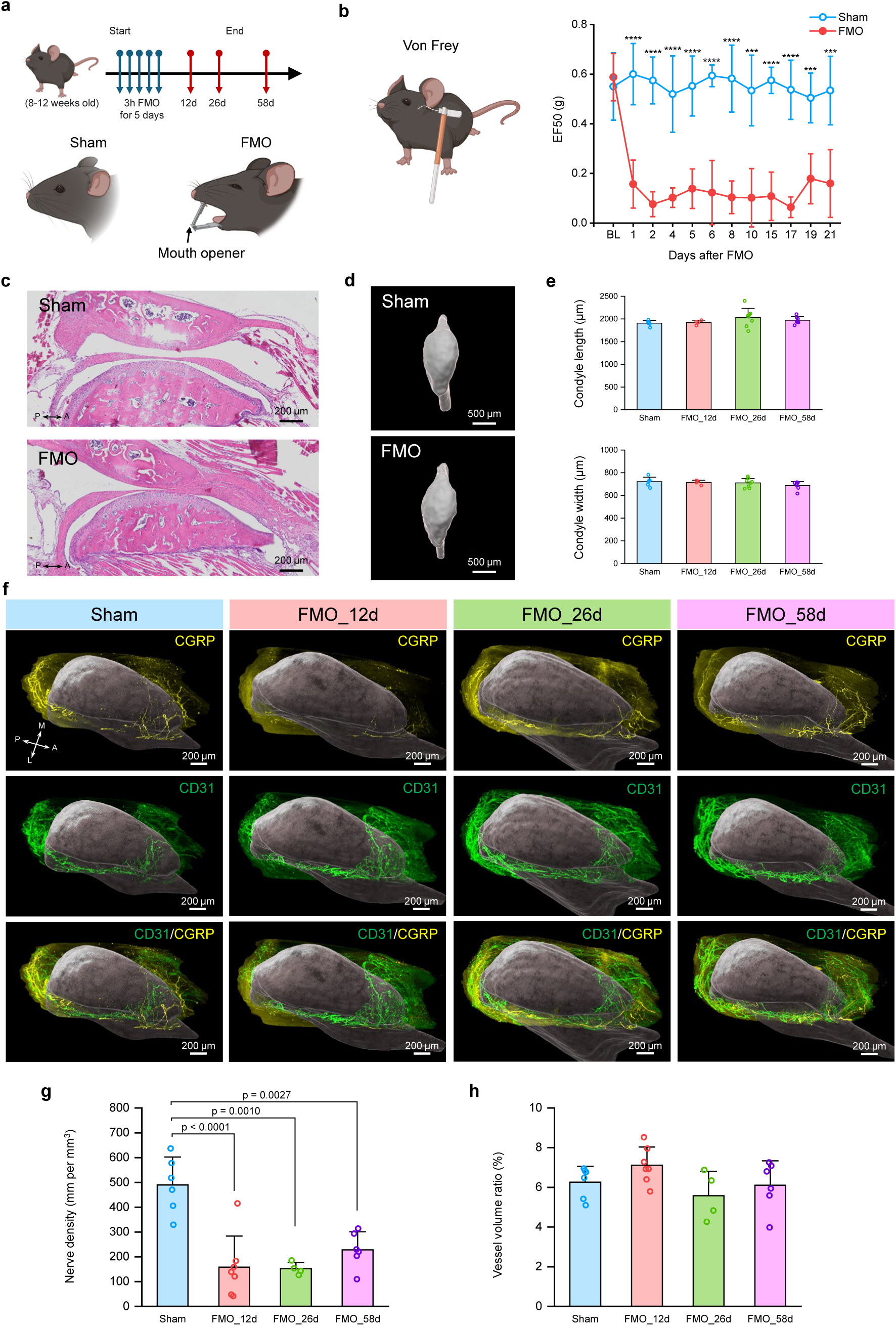
Whole joint neurovascular mapping in forced mouth opening (FMO) TMJ injury model. **a.** Schematic timeline for the FMO TMJ injury model. A customized mouth opener was applied between the maxillary and mandibular incisors to cause TMJ injury in the FMO group. **b.** Von Frey filament pain behavior testing and quantitative head withdrawal thresholds (EF50) in Sham and FMO mice over a period of 21 days (figure is reproduced from our published data in ref 34). ***p<0.001, ****p<0.0001. p-value was determined by a two-sided t-test. **c.** H&E staining results with mouse TMJs in Sham and FMO mice at 58 days after the FMO procedure. Scale bar, 200 µm. **d.** 3D reconstruction of mouse TMJ condyle heads in Sham and FMO mice at 12 days after FMO procedure. Scale bar, 500 µm. **e.** Quantitative condyle size measurements based on condyle µCT 3D reconstruction with Sham (n = 6 measurements from 6 mice), FMO_12d (n = 4 measurements from 4 mice), FMO_26d (n = 8 measurements from 8 mice), FMO_58d (n = 7 measurements from 7 mice). **f.** 3D reconstruction results of whole joint neurovascular mapping in mouse TMJ of Sham and FMO mice. Neurovascular data in the TMJ disc was only displayed for comparison. The TMJ condyles were segmented from the mapping data and rendered in grey to provide a spatial reference. Scale bar, 200 µm. **g.** Quantitative nerve density in Sham (n = 6 measurements from 4 mice), FMO_12d (n = 7 measurements from 5 mice), FMO_26d (n = 4 measurements from 2 mice), FMO_58d (n = 6 measurements from 4 mice). p-value was determined by one-way ANOVA with Bonferroni post-hoc test. **h.** Quantitative blood vessel density in Sham (n = 6 measurements from 4 mice), FMO_12d (n = 7 measurements from 5 mice), FMO_26d (n = 4 measurements from 2 mice), FMO_58d (n = 6 measurements from 4 mice). The nerve and vessel densities were quantified at the anterior region of the TMJ disc (**Supplementary Fig. 4d-4e**). All data depict mean ± standard deviation. A, anterior; P, posterior; M, medial; L, lateral.

### Whole joint neurovascular mapping in mouse knees

TMJs and knee joints are two major synovial joints in the body, sharing similarities and having intrinsic differences. To demonstrate the broad utility of our method, we applied the MUSIC protocol used for the mouse TMJs to mouse knees. We achieved successful tissue clearing of mouse knee joints (**Fig. 4a**) and labeled the pain-sensing CGRP-positive nerve fibers throughout the intact mouse knee (**Fig. 4b**). From 2D optical sections collected at different knee locations, we can clearly see a high density of CGRP nerve distributions in the fat pad, capsule, and meniscus peripheral tissues (**Fig. 4b**). Nerve fibers were also seen in the ligaments at the center of the knee, demonstrating the complete penetration of antibodies into the knee (**Fig. 4b**). Beyond a glimpse of the knee joint innervation pattern through separate 2D optical sections as done traditionally, we can now present the neural network in the entire knee through 3D reconstruction and trace the path of continuous long nerve fibers and their connectivity, as well as their spatial location and distributions within the joint anatomy (**Fig. 4c** and **Supplementary Fig. 3c-3d**). Zoomed to the lateral region of the knee (highlighted in a dashed red square), nerves branched out from a major nerve bundle and innervated the fat pad, meniscus peripheral tissues, and capsule (**Fig. 4c**). Moreover, we investigated how the neurovascular structure changes under degenerative conditions in the knee of the *Prg4^-/-^* mouse model. As reported in the literature, TMJs and knees developed osteoarthritis-like degenerative changes in *Prg4^-/-^* mice ^32,44^. Our results revealed an increased neurovascular density in the *Prg4^-/-^*mouse knee joint, confirming other studies ^32,44^ (**Fig. 4d** and **Supplementary Movie 10-11**). Benefiting from whole joint mapping, we can see how the neurovascular architectures change with dense nerves and vessels radiating through the *Prg4^-/-^*knee joint (**Fig. 4d**). Our results demonstrate the application of our method within the broader orthopedic field, opening new opportunities to investigate knee joint diseases.

**Figure 4.**
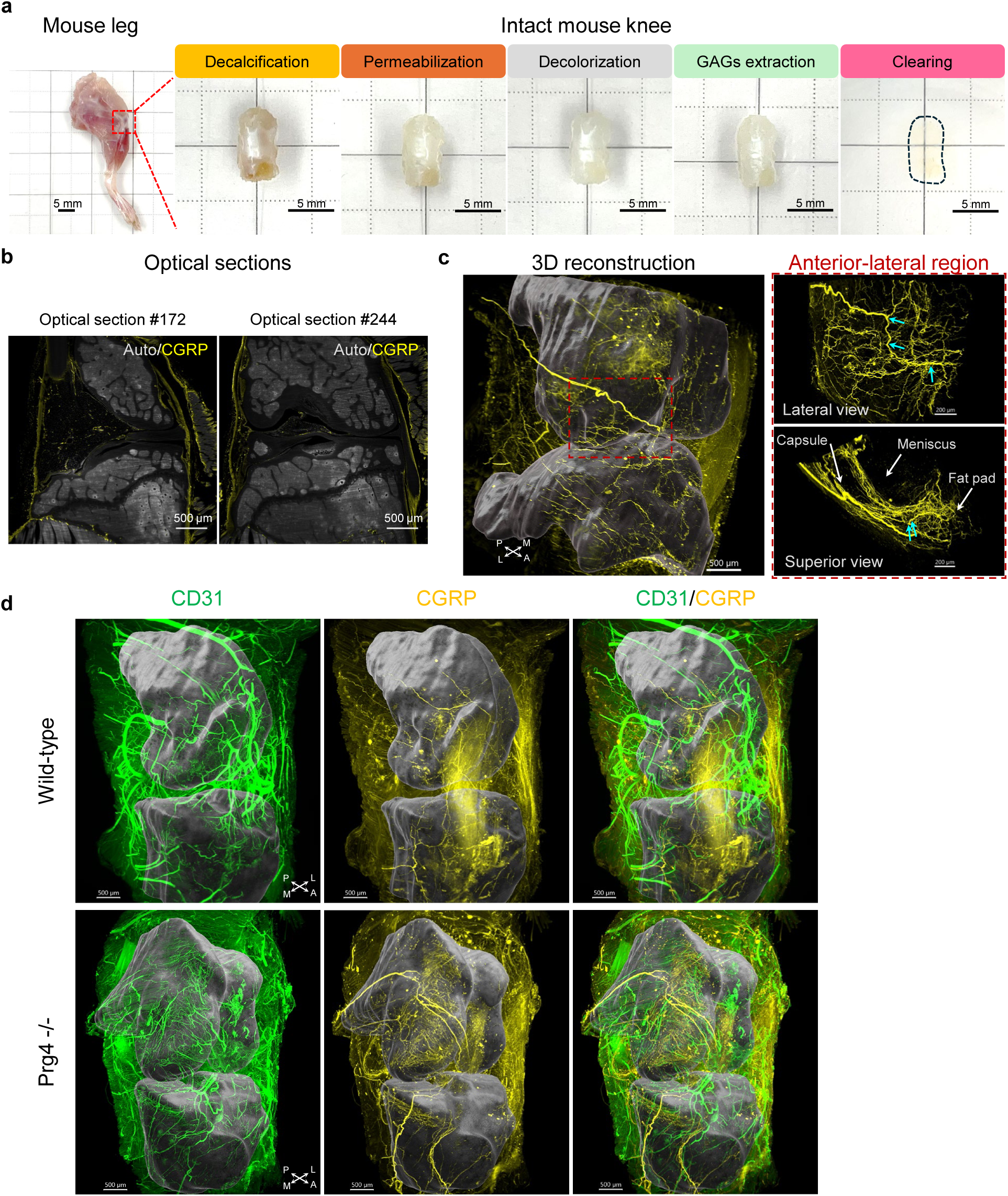
Whole joint neurovascular mapping in mouse knees. **a.** Photos of mouse leg and mouse knee along MUSIC procedures. The red dashed square highlighted the region where the mouse knee was harvested. The black dashed line depicted the contour of a cleared mouse knee. Scale bar, 5 mm. **b.** 2D optical sections from the 3D neural mapping in a mouse knee. Autofluorescence (Auto) signals were collected using a 488 nm laser and an emission filter of 525/50 nm. Scale bar, 500 µm. **c.** 3D reconstruction results of whole joint mapping in a mouse knee. Left, 3D whole joint reconstruction. The femur and tibia bones were segmented and rendered in dark grey to provide a spatial reference (**Supplementary Fig. 3c-3d**). The red dashed square highlighted the lateral region of the mouse knee for high-resolution imaging. Scale bar, 500 µm. Right, zoom-in views at the lateral region of the mouse knee. Cyan arrows showed the nerves branching points. The joint capsule, meniscus, and fat pad were indicated with a white arrow. Scale bar, 200 µm. **d.** 3D reconstruction results of whole joint mapping in mouse knees in wild-type and Prg4^-/-^ mice. The femur and tibia bones were segmented and rendered in dark grey to provide a spatial reference. Scale bar, 500 µm. A, anterior; P, posterior; M, medial; L, lateral.

### Scalable 3D mapping in joints of larger animal species

Various animal species have been used in musculoskeletal studies to pave the way for clinical research. Commonly used preclinical models include mice, rats, and pigs, with increasing joint dimensions as the animal size increases. 3D mapping of large-sized joints presents critical challenges due to tissue clearing and immunostaining difficulties. Scalability issues were well considered when our method was initially developed. We strategically used CHAPS and guanidine treatments for deep permeabilization; these two chemicals are proven to perform excellently for large human tissues ^31^. Sufficient joint permeabilization allows effective tissue homogenization and antibody penetration to facilitate clearing and immunostaining of large samples. In addition, using heptakis(2,6-di-O-methyl)-β-cyclodextrin minimizes the antibody aggregation issue during prolonged antibody incubation ^35^. We successfully cleared and mapped the rat TMJs (∼10 mm), knees (∼10 mm), and spine samples (∼7 mm), as well as pig TMJ discs with surrounding tissues (∼40 mm) (**Fig. 5a** and **Supplementary Fig. 6**), demonstrating the scalability and versatility of our method for musculoskeletal applications. Our results on rat TMJs showed densely distributed blood vessels in the anterior and posterior regions of the TMJ disc (**Fig. 5b** and **Supplementary Movie 12**), which agrees with the mouse data, supporting the conservation of neurovascular structures among species. Close-looped vessels found at the border of the disc vasculature verified the absence of vessels in the disc center region (**Fig. 5c**). Rat knee mapping data revealed a similar innervation pattern to the mouse knee, with high-density innervation in the joint capsule and meniscus peripheral tissues (**Fig. 5d**, **5e** and **Supplementary Movie 13**).

**Figure 5.**
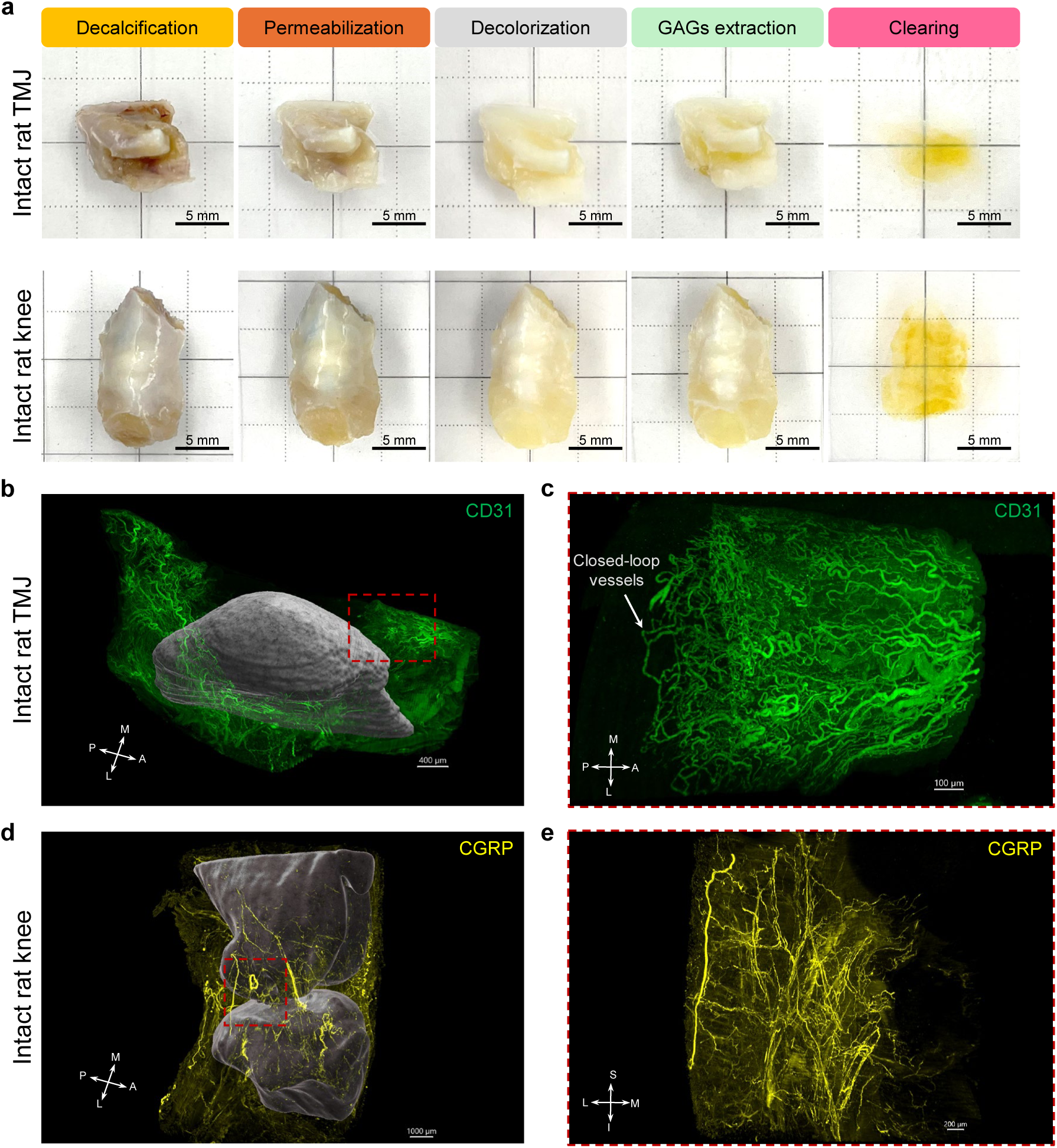
Scalable whole joint neurovascular mapping in large animal samples. **a.** Photos of rat TMJs and knees along MUSIC procedures. Scale bar, 5 mm. **b.** 3D reconstruction results of whole joint vascular mapping in a rat TMJ. TMJ condyle was segmented and rendered in grey to provide a spatial reference. The red dashed square depicted the anterior region of the rat TMJ disc for high-resolution imaging. Scale bar, 400 µm. **c.** Zoom-in views of the anterior region of the rat TMJ. The white arrow highlighted the closed-loop vessels at the edge of vascularization. Scale bar, 100 µm. **d.** 3D reconstruction results of whole joint neural mapping in a rat knee. Knee femur and tibia bones were segmented and rendered in dark grey to provide a spatial reference. The red dashed square depicted the lateral region of the rat knee for high-resolution imaging. Scale bar, 1000 µm. **e.** Zoom-in views of the lateral region of the rat knee. Scale bar, 200 µm. A, anterior; P, posterior; M, medial; L, lateral; S, superior; I, inferior.

In addition, we cleared and imaged the neural structure of the rat spine at the sacrum sections (**Supplementary Fig. 6a-6b**). High-resolution imaging data revealed the pain-sensing innervation at the intersection of two sacrum sections (**Supplementary Fig. 6c** and **Supplementary Movie 14**). Similarly, a large pig TMJ disc with surrounding tissues with a size of 33.5 mm × 43.8 mm × 15.6 mm was also successfully immunostained and cleared. No nerves were observed at the disc central region, with fiber bundles at the lateral, anterior, and posterior regions (**Supplementary Fig. 6d-6f** and **Supplementary Movie 15**). These results illustrated the scalability and versatility of our method, providing a platform method for musculoskeletal research.

## CONCLUSION

Understanding musculoskeletal joints from a multiscale perspective, from the molecular to the anatomical realm, is essential for resolving the confounding relationships between structure and pain, elucidating the intricate mechanisms regulating joint health and diseases, and guiding the development of new treatment strategies. In this study, we developed the MUSIC method for 3D whole joint cellular and molecular mapping by resolving key challenges faced with joint immunostaining and tissue clearing and integrating with advanced large-field light sheet imaging. Our method achieves simultaneous 3D imaging of both macroscale joint anatomy and microscale profiles of various joint structures (e.g., neurovascular networks) without the need for image coregistration. We demonstrated the power of our method by applying it to multiple joint systems (i.e., TMJs, knees, and spines) and various species (i.e., mice, rats, and pigs). Our method also allows comprehensive and systematic whole-joint investigation of microscale changes when the joint degenerates or gets injured, providing multiscale data critical for dissecting the relationship between joint structure and pain and studying whole joint health.

The musculoskeletal joint is a complex organ system composed of diverse tissue types, each exhibiting distinct hierarchical structures that collectively contribute to joint function, homeostasis, and disease progression ^4,42^. An integrated, multiscale approach is needed to investigate joint problems at the whole-joint level but with molecular-level details. Our MUSIC 3D whole joint mapping method provides a perfect tool to advance the field. Compared to the traditional 2D histological analysis method, 3D whole joint mapping with MUSIC offers significant advantages. First, it does not require sample sectioning, keeping the joint intact and maintaining the spatial relationships among joint components. Second, it captures all the features from a 3D and whole joint perspective, allowing non-skeptical and non-biased data interpretation and more accurate data quantification, particularly for structures like nerves and vessels with complex and heterogeneous 3D networks, as shown in our data. 3D mapping can resolve the controversial reports in the literature regarding the nerve density changes in diseased TMJs and knee joints in animal models ^43,51^. 3D mapping also collects multimodality data simultaneously, facilitating data registration and integration. Third, it allows better detection of “rare” events that are likely missed with 2D approaches, such as nerve branching from the main nerve bundle (**Fig. 1d** and **Fig. 4c**) and innervation at the intersection of sacrum segments (**Supplementary Fig. 6c**). Given the significance of neurovascular networks on joint health, disease development, and treatment outcomes, mapping their spatial organization provides a valuable opportunity to study how they contribute to joint structure integrity, function, and pain. We showed the neurovascular changes as joints degenerate and get injured, revealing the microscopic-level structural responses at different disease stages that complement the macroscopic-level changes, such as tissue morphology changes accessed through conventional imaging modalities (e.g., CT and MRI). We demonstrated that microscale changes in joint innervation, rather than macroscale joint structural alterations, are correlated with pain behaviors in FMO mice. These findings provide direct evidence linking joint microstructure to pain, highlighting a potential structural basis for pain sensation in joint disease. With the assistance of advanced computational modeling, we can even link the joint microstructure and macrostructure to joint biomechanics, joint function, and joint pain ^52^. Our mapping data contains anatomical and morphological information of hard and soft tissues and microscopic cellular and molecular information, which is the ideal dataset to establish an integrated computational modeling linking joint structure, function, and pain. Our data also provides insights into how each joint tissue contributes to different diseases. Our results showed different structural changes between the degenerative and injured models, highlighting the need for whole joint mapping to capture disease features. Moreover, our data on healthy wild-type animals provide a baseline, microscopic-level neurovascular map. The heterogeneity of neurovascular structures within a single tissue and across different tissues explains joint complexity, and the similarity of overall neurovascular distribution between mice and rats highlights joint conservation. Our data also uncovered the physical neurovascular connections among different joint components, which supports the whole joint concept and indicates that changes in one joint tissue might directly affect the other connected joint unit. Additionally, joint- mimicking in vitro models are new opportunities to understand joint diseases and develop/test drugs ^53^. Our data can contribute to this field by providing a joint template with spatial context. For instance, the neurovascular components may need to be considered in the design of the TMJ disc unit, given their presence at the peripheral disc shown in our results.

Tissue clearing and immunostaining were used in our method to achieve 3D whole joint mapping. Tissue clearing in the musculoskeletal field is in its infancy. Only a handful of tissue-clearing protocols have been developed in the literature for single-type mouse joint tissues ^20–25^. To date, only one very recently published protocol has reported whole-joint clearing of the mouse knee with the use of collagenase, a method that carries the risk of compromising tissue structure ^54^, and no studies have demonstrated its application to diverse tissue structures in various joint types across different species. Although the clearing of the entire mouse body has recently been reported, these studies are not focused on the joints ^35,55^. Their performance on the joints has yet to be demonstrated, and their application is also limited to mice due to the high cost of whole-body clearing and labeling for large animals. Our method specifically addresses this challenging task based on the unique structure and composition features of musculoskeletal joints and achieves a broad application in different types of joints across multiple animal species. Due to the limited availability of fresh human joint specimens, the application of our method in human samples has yet to be demonstrated here. Since the size of pig TMJ samples used in this study is much larger than adult human TMJ samples, we do not anticipate major issues applying our methods to human TMJ samples. The performance of our method on large human knees and spines will need further investigation. Moreover, utilizing immunostaining as the labeling approach in our method provides flexibility and opportunities to study joint cellular and molecular structures other than neurovascular structures.

3D imaging is becoming increasingly popular, given the tremendous, exciting information it can provide. Many 3D imaging platforms have been developed. Light sheet microscopes stand out with their capacity for large field of view and high-speed imaging as both benefit whole joint mapping. Open-sourced and low-cost light sheet microscopes have been developed over the years ^56–59^. 3D whole joint mapping using our method is anticipated to become a routine research tool to boost future research advancements. Additionally, other multiplexed imaging modalities, such as fluorescence lifetime imaging ^60^ and Raman spectroscopy ^61,62^, can be integrated with our method to achieve whole joint mapping of various subtype cellular and molecular structures. Due to the complexity of the joint and proximity of joint components, our data analysis is primarily based on manual or semiautomatic, labor-intensive methods for segmentation (hours to days of labor work for each dataset). Future research on developing artificial intelligence-based automatic segmentation methods suitable for the whole joint mapping dataset is critical to streamlining the entire procedure and extracting more insightful and quantitative information.

With the increased accessibility of 3D imaging systems and the advancement of data analysis methods, our method will deepen our understanding of joint health and disease mechanisms and contribute to a new and effective treatment strategy.

## EXPERIMENTAL SECTION

### Sample preparation

#### *Prg4^-/-^* mouse samples

Adult wild-type and *Prg4^-/-^* C57BL/6 male or female mice (#025737, Jackson Laboratory, Bar Harbor, ME) aged between 6-12 months old were utilized in this study to investigate the neurovascular patterns in joints with apparent osteoarthritis-like joint degeneration. Mice usage was approved by the Institutional Animal Care and Use Committee (IACUC) at Columbia University Irving Medical Center (AC-AABP1553). After euthanasia, mouse skins were carefully dissected from the whole body without damaging major blood vessels and causing bleeding. Then, the whole mouse body was placed in 10% formalin for fixation at 4°C for one day. This blood retention procedure is intended to retain as much blood inside the vessels as possible to enhance its fluorescence intensity with blood autofluorescence. Afterward, TMJs and knees were dissected and incubated in 10% formalin solution at 4°C for another two days to ensure thorough joint fixation. Fixed TMJs and knees were washed in 1x phosphate buffer saline (PBS) at least three times before further processing.

### Forced mouth opening (FMO) mouse samples

Adult C57BL/6 male or female mice (#000664, Jackson Laboratory, Bar Harbor, ME) aged between 8-12 weeks were used in this study with approval from the IACUC at the University of Maryland. The mice were anesthetized with isoflurane inhalation (3% for induction and 1.5% for maintenance) and kept in the isoflurane chamber during the procedure described in our previous protocol ^34^. FMO mice were injured with a customized mouth opener (0.017 in × 0.025 in rectangular wire) placed between the maxillary and mandibular incisors (**Fig. 3a**) for three hours per day for five consecutive days, while the Sham mice received no procedures. Following each procedure, both FMO and Sham mice were fed a soft diet (DietGel Recovery, ClearH2O, Westbrook, ME) and water gel (HydroGel, ClearH2O, Westbrook, ME). FMO mice were euthanized by transcardial perfusion with heparinized PBS followed by 4% paraformaldehyde in PBS at 12 days, 26 days, and 58 days after the final FMO procedure, respectively. Mice were randomly assigned to different groups. Sham mice were terminated at 12 days as reference control. Mouse heads were then harvested and washed with PBS three times for further processing.

### Rat samples

Fresh rat joints were collected from Sprague Dawley rats at the conclusion of other IACUC-approved research projects at the Medical University of South Carolina and Columbia University Irving Medical Center (AC-AABG8555). Rats were terminated and perfused transcardially with heparinized PBS and 4% paraformaldehyde in PBS. Rat TMJs were harvested from adult male or female rats aged 25-26 weeks, while rat knees and spines were obtained from adult female rats aged 10-20 weeks. Rat joints were then fixed in 4% paraformaldehyde for one day and washed in PBS.

### Pig samples

Fresh porcine heads (Yorkshire, 6 months) were obtained from a local abattoir and used within 12 h post-mortem. The pig TMJ disc with attached surrounding tissues was carefully dissected from the pig head and immediately fixed in 10% formalin solution at 4°C for three days and then washed with PBS three times before further processing.

### MUSIC protocol

As the joints have bony components, fixed joints were first decalcified with 20% EDTA- 2Na (dissolved in diH_2_O, pH = 8.0) (BDH4616, VWR, Radnor, PA) for 3-10 days with daily solution change at 37°C on a shaker. Complete joint decalcification was examined and confirmed with µCT imaging, showing uniform contrast throughout the joints. Then, joints were washed with 1x PBS three times and stored in PBS with 0.01% sodium azide at 4°C until further processing. Decalcified joints were incubated in 10% CHAPS (10232, Cepham Life Sciences, Fulton, MD) in diH_2_O at 37°C with gentle shaking for 7 days to deeply permeabilize the joint tissues. Joints were then dehydrated with graded methanol (154903, Sigma, St. Louis, MO) solutions (25%, 50%, 75%, 100%, 100%, mixed with diH_2_O) followed by delipidation treatment of dichloromethane (270997, Sigma, St. Louis, MO)/methanol mixture (v/v, 2:1) for 1-3 days until joints sink. Then, joints were bleached in chilled fresh 5% H_2_O_2_ (H1009, Sigma, St. Louis, MO) in methanol overnight at 4°C and rehydrated with methanol series (75%, 50%, 25%, PBS) at room temperature (RT). Joints were washed in Triton X-100 (T8787, Sigma, St. Louis, MO) /PBS mixture (v/v, 0.2% in 1x PBS) two times at RT. To loosen the joint extracellular matrix without significantly sacrificing the joint integrity, joints were incubated in a guanidine solution composed of 4M guanidine hydrochloride (97061, VWR, Radnor, PA), 0.05M sodium acetate (S7670, Sigma, St. Louis, MO) and 2% (w/v) Triton X-100 in 1x PBS (pH = 6.0) at RT for 3 days to remove the proteoglycans while keeping the collagen backbones. Then, the permeabilized joints were blocked in a blocking solution containing 6% donkey serum (017-000-121, Jackson ImmunoResearch, West Grove, PA), 10% dimethylsulfoxide (DMSO, BDH1115, VWR, Radnor, PA), and 0.2% Triton X-100 at 37°C for 3 days, and incubated in primary antibodies (**Supplementary Table 1** for a full list of antibodies used in this study) in an immunostaining buffer solution containing 3% donkey serum, 10% CHAPS, 2% Triton X-100, 10%DMSO, 1% glycine (G8898, Sigma, St. Louis, MO), 1% heptakis(2,6-di-O-methyl)-β-cyclodextrin (77158, VWR, Radnor, PA) in 1x PBS at 37°C for 3-7 days depending on the sample size. Joints were then washed 3-5 times in 1x PBS at 37°C for 3 days followed by incubation in an immunostaining buffer solution with secondary Alexa fluorescent dye-conjugated secondary antibodies, including Alexa Fluor 647 donkey anti-goat IgG antibody (A-21447, ThermoFisher, Waltham, MA), Alexa Fluor 647 donkey anti-mouse IgG antibody (A-31571, ThermoFisher, Waltham, MA), and Alexa Fluor 594 donkey anti-rabbit IgG antibody (A-31573, ThermoFisher, Waltham, MA) at 37°C for 3-7 days. All secondary antibodies used 1:500 dilution. Joints were washed 3-5 times in 1x PBS at 37°C for 3 days. All immunostaining steps were performed with gentle shaking to facilitate antibody penetration and washout. After the completion of immunostaining, joints were dehydrated in methanol series at RT for one day, delipidated in 100% DCM solution at RT until joints sink, and cleared in dibenzyl ether (DBE, 108014, Sigma, St. Louis, MO) at RT for 1-2 days until joints became optically transparent (**Fig. 1b**, **4a**, **5a**, and **Supplementary Fig. 6**). Detailed protocols for different joints and species are available in **Supplementary Table 2**.

### Light sheet imaging

Cleared whole joint samples were transferred to a non-toxic ethyl-3-phenylprop-2-enoate (ECi, 76806, VWR, Radnor, PA) solution with a similar refractive index to DBE prior to imaging^63^. Mice and rat joints were imaged with an Ultramicroscope II (Miltenyi Biotec, Germany) with a Super Plan configuration, which is equipped with a sCMOS camera (4.2 Megapixel, Andor Technology, UK), five excitation lasers (405 nm, 488 nm, 561 nm, 639 nm, 785 nm), tube lenses for post-magnification adjustment (0.6x, 1x, 1.67x, and 2.5x), and three objective lenses with dipping caps specifically designed and optimized for large-field light sheet imaging (1.1x/NA 0.1, 4x/NA 0.35, and 12x/NA 0.53). Cleared joints were mounted on the sample holder and immersed in the cuvette filled with ECi solution. 3D image stacks were acquired using the following laser and filter settings: 488 nm laser, emission 525/50 nm for autofluorescence signal; 561 nm laser, emission 595/40 nm for fluorescence signals from Alexa Fluor 594 dye; 639 nm laser, emission 680/30 nm for fluorescence signals from Alexa Fluor 647 dye. Whole joint mapping with mouse TMJs was imaged with the 4x objective at 1x magnification at a z-step interval of 1.62 µm, matched with the in-plane pixel dimension. In contrast, mouse knees, rat TMJs, and rat spines were imaged with the 4x objective at 0.6x magnification at a z-step interval of 2.71 µm. Rat knees were imaged with the 1.1x objective at 1x magnification at a z-step interval of 5-20 µm. 3D data acquisition in the regions of interest were collected with a 4x objective at 1.66x magnification or a 12x objective at 1x magnification at a z-step interval of 1 µm.

Pig TMJ samples were imaged with an UltraMicroscope Blaze (Miltenyi Biotec, Germany) equipped with a sCMOS camera (4.2 Megapixel, Andor Technology, UK) and a special objective lens with dipping caps. Pig TMJ samples were also imaged in ECi solution. Mosaic acquisition was performed with a 4×2 tile scan at a 10% overlap to cover the entire sample. The pig TMJ was imaged with the 1.1x objective at 1x magnification at a z-step interval of 8 µm throughout the full thickness of the sample. 3D image stacks were acquired using a 639 nm laser and an emission filter of 680/30 nm.

### 3D mapping data reconstruction and quantification

3D image stacks were saved in TIFF format for each channel separately. ImarisFile Converter (version 9.9, Bitplane, UK) was used to convert the 3D image data into Imaris file format for further data processing. The tile scan data with pig TMJ was stitched with Imaris Stitcher (version 9.9, Bitplane, UK). The decalcified bone region, soft tissues (e.g., cartilage and meniscus), and muscles were visualized using a 561 nm laser and an emission filter of 595/40 nm (**Supplementary Fig. 3a, 3c**). Each joint tissue was manually segmented using the “Surface” function in Imaris, and a surface was eventually created to reconstruct its 3D geometry (**Supplementary Fig. 3b, 3d**). The segmented surfaces were rendered with different colors and textures in Imaris for visualization. The reconstructed joint structure can be used as a mask to highlight the neurovascular structure of each joint tissue. Due to the relatively high autofluorescence from the muscles, they were not displayed in all the TMJ data for better visualization of the joint neurovascular structures. Vessels and nerves within the mouse TMJ disc were only displayed in the FMO data from the whole joint mapping (**Fig. 3f**). Noise signals in the mapping data due to the presence of particles and antibody entrapments were removed based on their unique features of high brightness and spheric shape.

The neurovascular structure was quantified in the lateral capsule with the synovial membrane of the TMJ in the *Prg4^-/-^* data and the anterior region of the TMJ disc in the FMO data. For the quantification in *Prg4^-/-^*data, the TMJ lateral capsule with synovial membrane at the superior joint cavity was manually segmented near the intersection of the zygomaticomandibularis muscle and zygomatic arch with a thickness of 100 µm (**Supplementary Fig. 4a**). For the quantification of FMO data, a region of interest with a size of 300 × 300 × 243 µm^3^ (185 × 185 × 150 voxels) was used to crop the anterior region of the TMJ disc (**Supplementary Fig. 4d**). The location of the anterior TMJ disc was identified with reference to the underneath TMJ condyle head. Nerve fibers were traced manually using the “Filament” function and the “AutoPath” method, while blood vessels were automatically segmented using the “Surface” function (**Supplementary Fig. 4c, 4e**). The selected capsule and disc volume was then quantified using the “Surface” function and automatic segmentation. The total length of the nerve fibers and the total volume of blood vessels were recorded and normalized to the disc volume to calculate the nerve and vessel density.

### Histology analysis

Fixed mice TMJs were decalcified and prepared for paraffin or frozen embedded sections. Sagittal sections of the TMJ were collected and stained with Hematoxylin & Eosin (H&E). Bright- field images were captured on a Keyence BZ-X800E microscope (Keyence, Raleigh, NC).

### Micro-computed tomography (µCT) imaging and quantification

Fixed mice TMJs from the *Prg4^-/-^* and FMO models were scanned with a Scanco µCT 40 device (Scanco Medical, Southeastern, PA) at a voxel resolution of 18 µm to capture the bone morphology. The µCT dataset of the bony structure of TMJs was then imported to Imaris (version 10.1, Bitplane, UK). The “Surface” function of Imaris was used to reconstruct the TMJ condyle head. According to the literature definition ^64^, the length and width of the condyle head were then determined by measuring the distance between the outermost points of the anterior, posterior, lateral, and medial of the condyle, respectively, using the “Measurement Points” function of Imaris.

### Pain behavior assessment using the von Frey assay

The von Frey (VF) test using the up-down technique was utilized to assess changes in mechanical sensitivity, as used in our prior publication ^34^. The orofacial skin over the TMJ was exposed to VF filaments with bending forces of 0.008 g, 0.02 g, 0.04 g, 0.07 g, 0.16 g, 0.4 g, 0.6 g, 1.0 g, 1.4 g, and 2 g, which were calibrated using an electronic balance. The nocifensive reaction was described as rapid or vigorous head withdrawal from the probing filament. Each VF filament was applied 5 times, each lasting a few seconds. The response frequencies [(the number of responses/the number of stimuli) × 100%] to a range of VF filament forces were determined, and S-R curves were plotted. Mechanical thresholds were measured as the force necessary to induce a 50% response frequency (EF50).

### Statistical analysis

A two-sided t-test was performed to evaluate the condyle head length and width differences and the nerve fiber density and blood vessel density differences between the wild-type and knockout groups in the *Prg4^-/-^* model and the withdraw threshold differences in von Frey test at different days after FMO between the Sham and FMO groups. One-way ANOVA with Bonferroni post-hoc test was used to examine the condyle head length and width differences and the nerve fiber density and blood vessel density differences among the experimental groups in the FMO model. The animal number was taken as a random factor for the analysis. All analyses were performed in SPSS (IBM SPSS Statistics, Version 28.0, IBM Corp., Armonk, NY). Significant differences were reported at p<0.05, with descriptive statistics reported as mean ± standard deviation.

## Data availability

Raw image data of this study are available from the corresponding author upon reasonable request.

## Acknowledgments

This work was supported by the National Institutes of Health (NIH) grants P20GM121342 and R01DE021134 to H.Y.; R34DE033593 and U01DE031512 to H.Y. and J.S.L.; R21GM104683 and P20GM103499 to T.Y.; R01DE031477 to M.K.C. and Y.S.K., and R35DE030045 to M.K.C.; R01DE029068 to M.C.E..

We would like to thank Dr. Alexander Gruhl and JD Shipp from Miltenyi Biotec for their help with the pig TMJ imaging using the UltraMicroscope Blaze. We also want to thank Isabelle Museck, Alex Levin, Bo Suthon, and Saadhana Arunkumar for their help with image segmentation and Dr. Wenyu Gou from the Medical University of South Carolina for her help with the animal sample collection. Schematic figures were created on the Biorender website (https://www.biorender.com/).

## Author contributions

The manuscript was written with contributions from all authors. All authors have given approval to the final version of the manuscript. H.Y. conceived and directed the study. H.Y., T.Y., P.C., and J.C. developed the method for tissue clearing and imaging. H.Y., P.C., T.Y., M.K.C., M.C.E., and J.S.L. designed the experiments. P.C. and J.C. performed all the joint clearing and imaging experiments. P.C., J.C., A.S., R.G.H., B.K., I.A., S.R., I.T., B.J.D., S.W. and Y.S.K. performed the µCT imaging, joint histology, quantitative measurements, and behavior testing. P.C., B.K., J.H., I.T., J.T.H., and M.Y. prepared the animals. P.C., J.C., and H.Y. analyzed and interpreted the mapping data. P.C., J.C., and H.Y. drafted the manuscript. P.C., J.C., M.K.C., M.C.E., J.S.L., T.Y., and H.Y. critically reviewed and edited the manuscript.

