## Supplementary Materials for "Multiscale 3D whole joint cellular and molecular mapping dissects the relationship between structure and pain"

**Content**

Supplementary Figures 1-6

Supplementary Tables 1-2

Supplementary Notes 1-2

Supplementary Movies 1-15

**Supplementary Figures:**

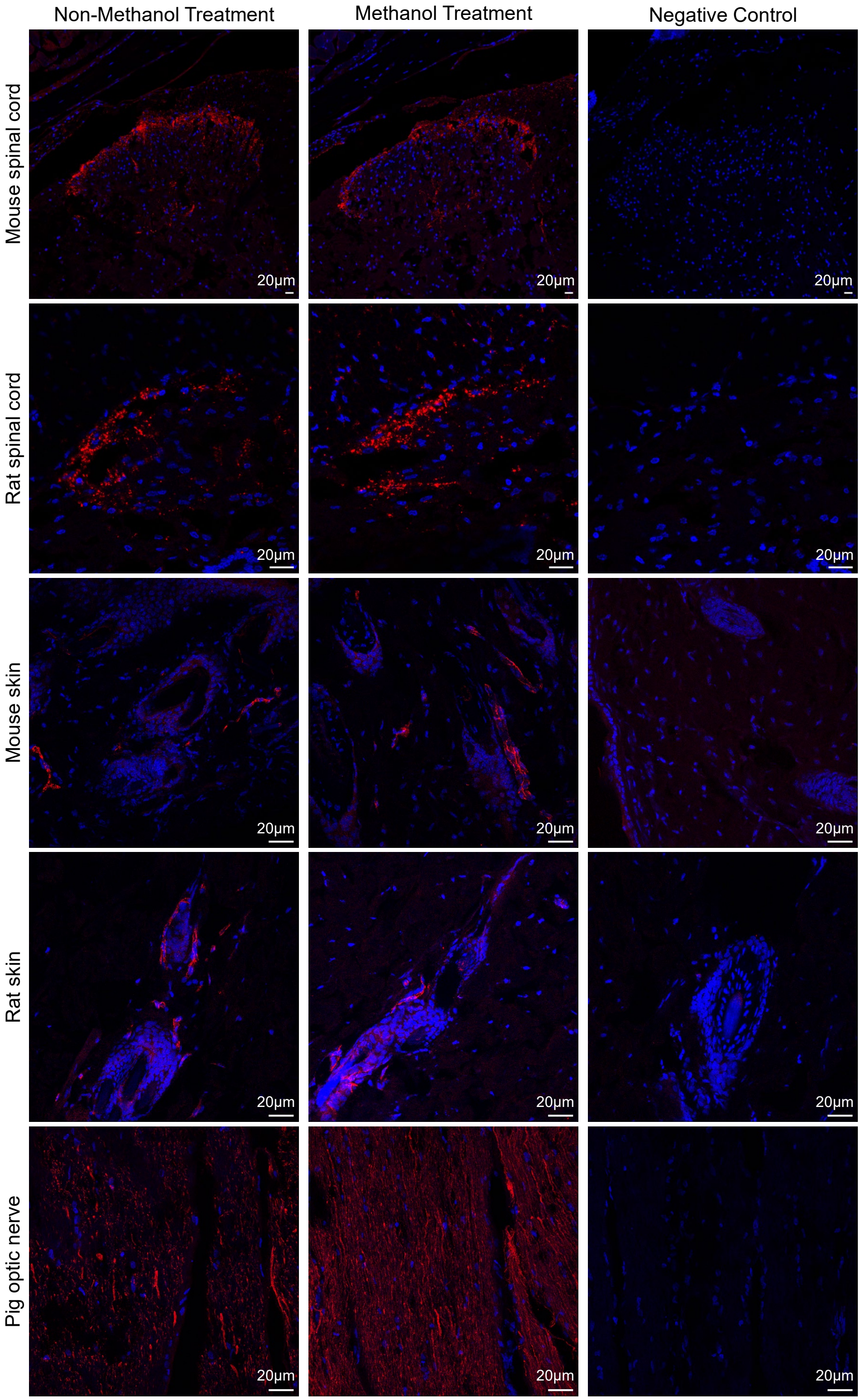


**Supplementary Figure 1. Antibody verification and compatibility with musculoskeletal joint immunostaining and clearing technique (MUSIC).** The Goat anti-CGRP antibody was used to stain the frozen sections of the mouse spinal cord (see antibody information and immunostaining methods in Supplementary Table 1 and Supplementary Note 1). Our results showed confined CGRP immunostaining at the outer lamina of the spinal dorsal horn, agreeing with the literature reports. Methanol treatment does not affect the fluorescence intensity, validating the compatibility of this antibody for the MUSIC method. Mouse anti-CGRP antibody was applied to label the rat spinal cord frozen section. Similar results were seen with CGRP staining at the surface of the spinal dorsal horn, and no apparent fluorescence changes were observed after methanol treatment. Rabbit anti-CD31 antibody was utilized to stain the mouse and rat skin. Vessels were observed in mouse and rat skin samples, and the same structures with similar fluorescence intensity were seen in the methanol treatment sections. Mouse anti-neurofilament 200 antibody was used to label pig optic nerve sections. Long nerve fibers running parallel to the optic nerve bundles were captured, and their fluorescence intensity was not affected after methanol treatment. All negative control samples showed no immunostaining signals. These results verified the specificity of the antibodies for the specific molecular targets and demonstrated their compatibility with the MUSIC method. Scale bar, 20 μm.

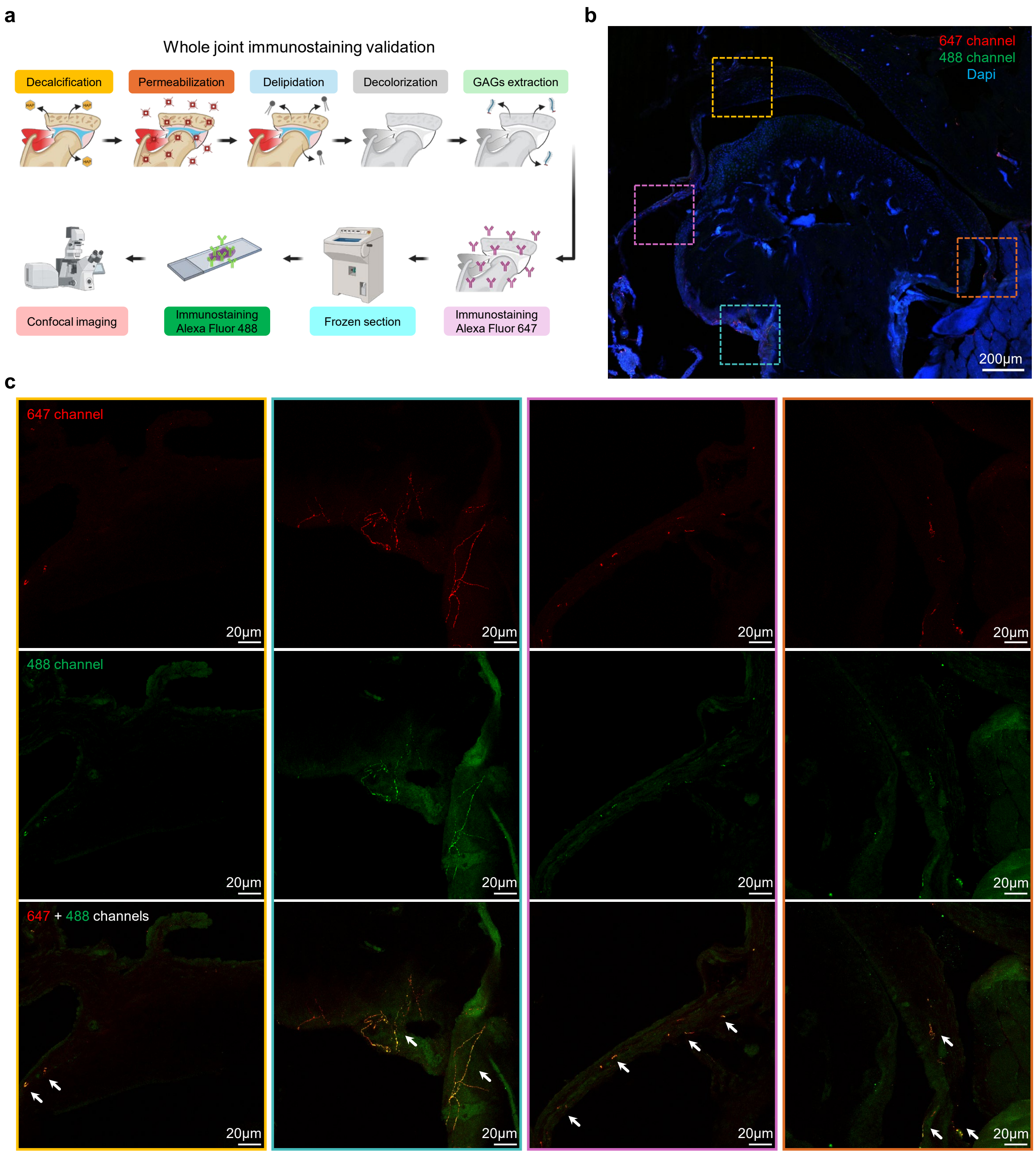


**Supplementary Figure 2. Whole joint immunostaining validation. a.** Schematic of sample preparation for whole joint immunostaining validation (see Supplementary Note 2). Mouse TMJs were processed following the MUSIC protocol for decalcification, permeabilization, delipidation, decolorization, and GAGs extraction. Then, whole TMJs were immunostained with primary antibodies for CGRP+ nerve fibers and secondary antibodies tagged with an Alexa Fluor 647 dye. Next, the immunostained joint was prepared for frozen section and cut into 10-15 µm sections and stained with the same primary antibody and a different secondary antibody with Alexa Fluor 488 dye. Joint sections were then imaged with a confocal microscope. **b.** A typical mouse TMJ section captured under the confocal microscope with a 20x objective. The dashed boxes highlight the regions for higher magnification imaging using a 40x objective. Scale bar, 20 μm. **c.** High-resolution imaging at the highlighted regions in **b**. Nerve fibers were detected in both channels, and their signals overlapped (white arrows), demonstrating that the whole joint immunostaining successfully labeled the target structures. Scale bar, 20 μm.

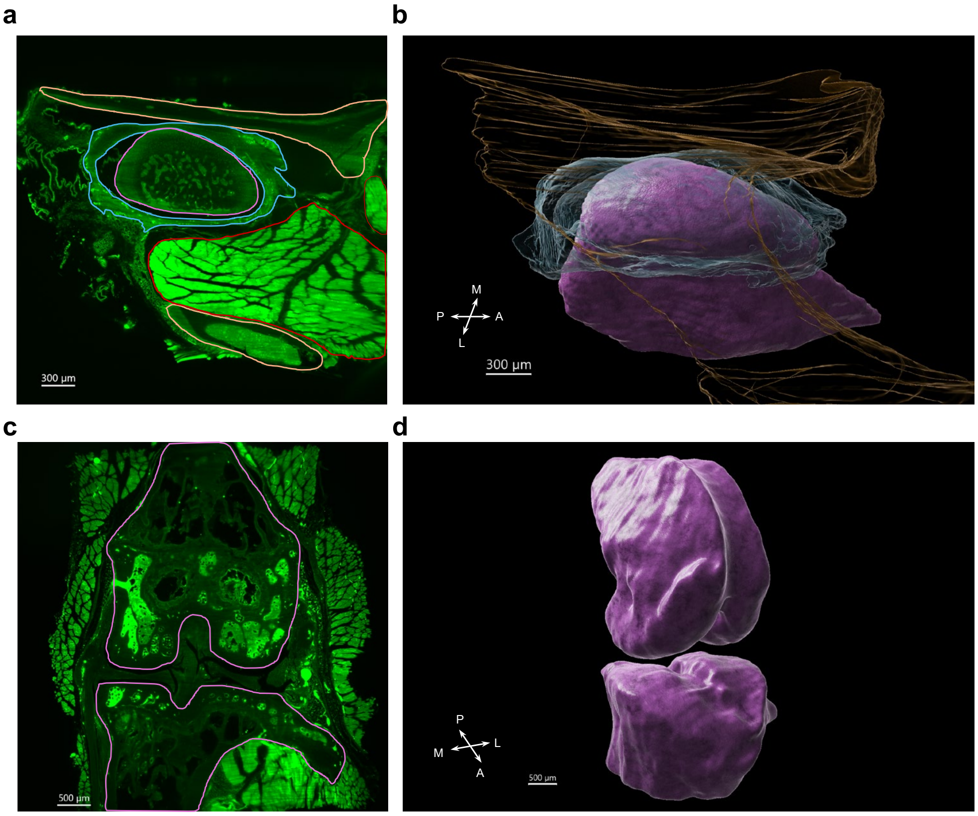


**Supplementary Figure 3. Joint mapping data and segmentation. a.** Typical 2D optical section collected from the mouse TMJ mapping data. The image was acquired using a 561 nm laser and an emission filter of 595/40 nm. The image intensity was intentionally enhanced for visualization. Each joint component was manually segmented in Imaris using the "Surface" function. The TMJ condyle with bone and cartilage is circled in purple, the disc is highlighted in cyan, the fossa is shown in orange, and the muscle is depicted in red. A series of 2D optical sections from the whole joint mapping dataset were segmented to reconstruct the 3D geometry of joint components. Scale bar, 300 μm. **b.** 3D reconstruction of major joint components in mouse TMJ. The TMJ condyle, disc, and fossa are shown in the colors of purple, cyan, and orange, respectively. The 3D surface of joint components was rendered in Imaris for visualization. Scale bar, 300 μm. **c.** Typical 2D optical section in the mouse TMJ mapping data. The image was acquired using the same settings in **a**. The femur and tibia bones, including the cartilage, are circled with purple lines. Scale bar, 500 μm. **d.** 3D reconstruction of femur and tibia in mouse knees. Scale bar, 500 μm. A, anterior; P, posterior; M, medial; L, lateral.

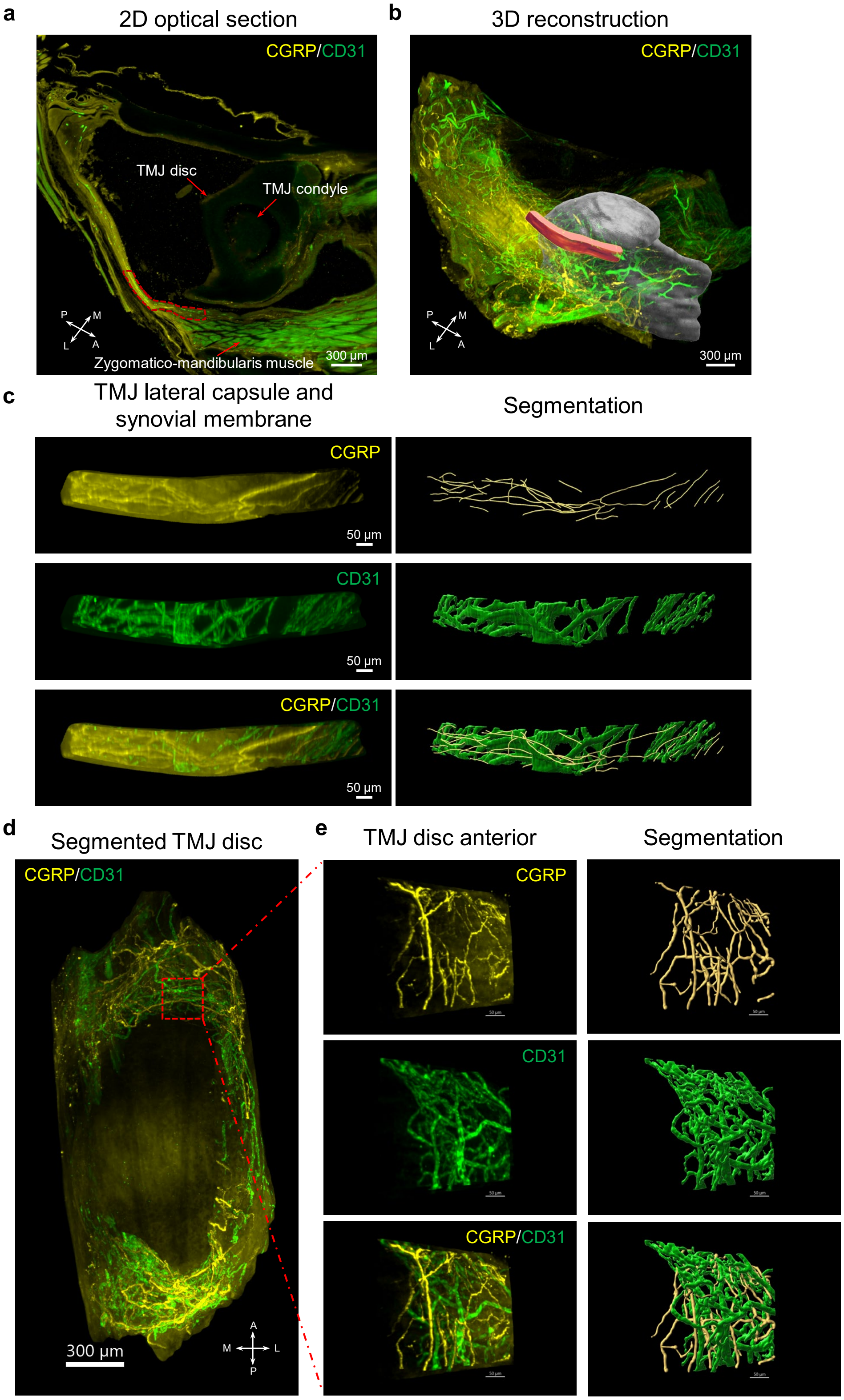


**Supplementary Figure 4. Joint mapping neurovascular structure quantification. a.** 2D optical section from the 3D neurovascular mapping in the whole mouse TMJ in *Prg4*$^{-/-}$ mice. The red dashed line highlights the segmented regions of the lateral capsule with the synovial membrane. Red arrows indicate the TMJ condyle, disc, and zygomatico-mandibularis muscles. **b.** 3D reconstruction of the 3D neurovascular mapping in the whole mouse TMJ in *Prg4*$^{-/-}$ mice. The 3D volume rendered in red color highlights the segmented regions of the lateral capsule with the synovial membrane. **c.** The nerve and blood vessel densities in the lateral capsule and synovial membrane region were quantified by calculating the nerve length and blood vessel volume in their corresponding channel. **d**. Neurovascular structure quantification in mouse TMJ disc. Neurovascular structure in mouse TMJ disc was masked with the disc surface geometry shown in Supplementary Fig. 3b. Scale bar, 300 μm. **e**. A region of interest at the anterior region of the TMJ disc (red dashed square in **d**) was cropped out, and the nerve and blood vessel densities were quantified by calculating the nerve length and blood vessel volume in their corresponding channel. Scale bar, 50 μm. A, anterior; P, posterior; M, medial; L, lateral.

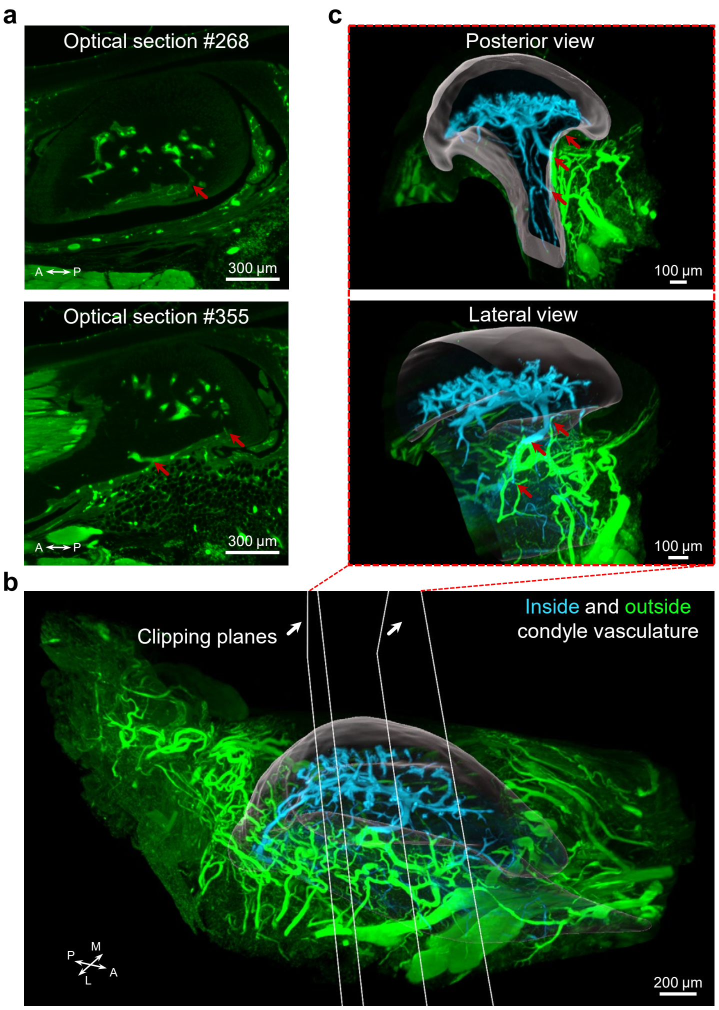


**Supplementary Figure 5. Vasculature connectivity inside and outside the TMJ condyle. a.** Optical sections of 3D vasculature mapping data in a mouse TMJ. The vasculature channels that connect the inside and outside of the condyle are highlighted with red arrows. Scale bar, 300 µm. **b.** 3D reconstruction of the 3D vasculature structures in mouse TMJ. The vasculature inside and outside the condyle is labeled as cyan and green colors, respectively. The condyle was segmented and rendered as a transparent grey color. The clipping planes were added to crop the middle sections of the 3D dataset to show a zoom-in view of the vasculature connectivity, as shown in **c**. Scale bar, 200 µm. **c.** Posterior and lateral view of the cropped sections of the 3D whole TMJ vasculature dataset. Red arrows highlight the connectivity point between the inside and outside condyle vasculature. Scale bar, 100 µm. A, anterior; P, posterior; M, medial; L, lateral.

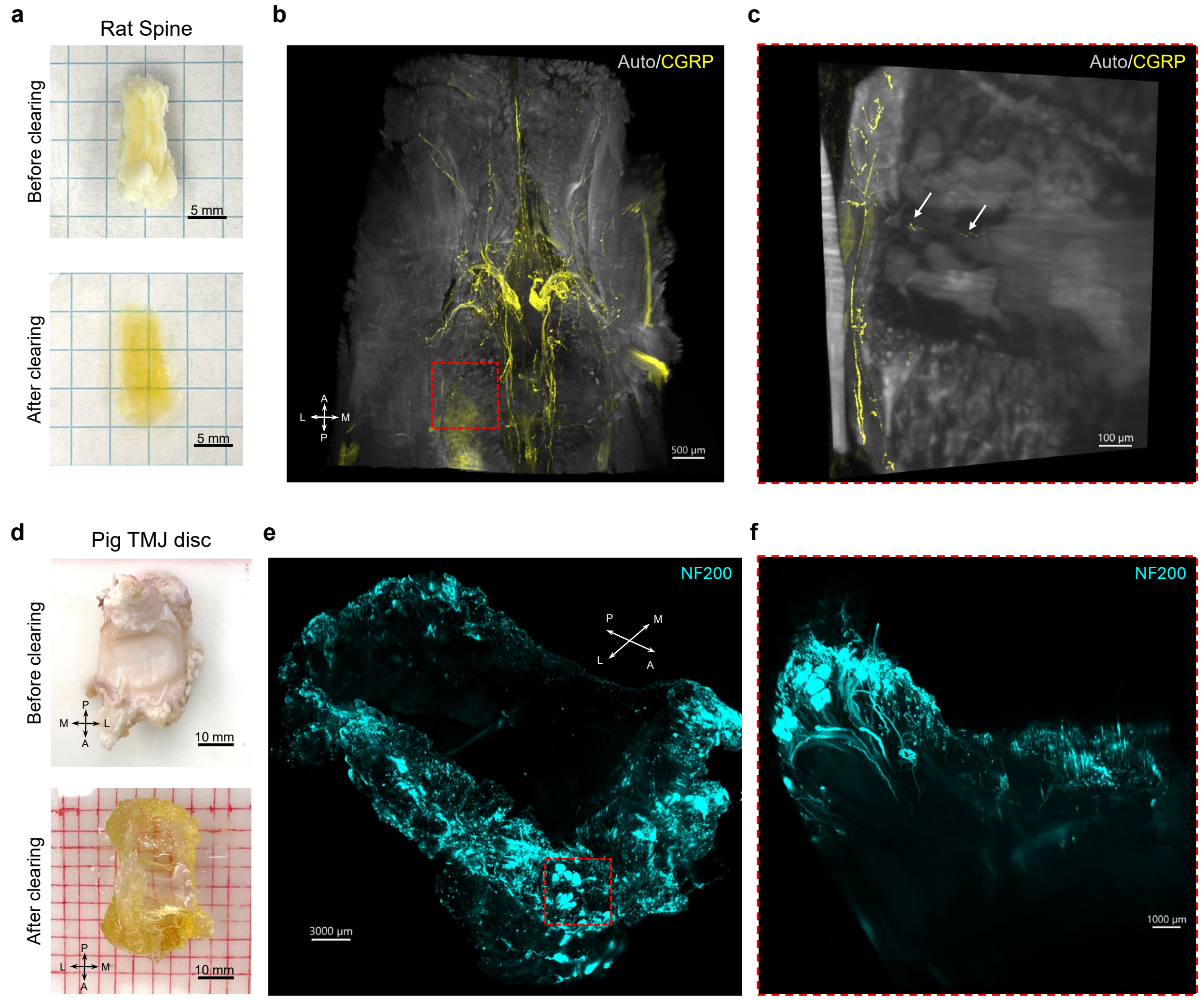


**Supplementary Figure 6. Whole joint mapping in large joint samples. a.** Photos with rat spine samples before and after tissue clearing. Scale bar, 5 mm. **b.** Whole joint mapping results with CGRP stained rat spine near the sacrum sections. CGRP signals were captured using a 639 nm laser with an emission setting of 680/30 nm. Autofluorescence (Auto) was collected using a 488 nm laser with an emission filter of 525/50 nm. Scale bar, 500 µm. **c.** High magnification mapping data at the intersection of two sacrum vertebrae, dashed red square shown in **b**. CGRP nerve fibers (arrows) were seen innervating the bony region between two sacrum vertebrae. Scale bar, 100 µm. **d.** Photos with pig TMJ disc samples before and after tissue clearing. This pig sample also includes the surrounding soft tissues, such as the retrodiscal tissue. The sample size is approximately 33.5 mm × 43.8 mm × 15.6 mm. Scale bar, 10 mm. **e.** The neural mapping in the entire pig TMJ immunostained with NF200. The high density of neural staining was observed in the anterior, lateral, and posterior regions. Scale bar, 3000 µm. **f.** High magnification mapping data at the anterior-lateral region as highlighted in **e** with a dashed red square. Large nerve bundles and small nerve structures were detected. Scale bar, 1000 µm

**Supplementary Tables:**

**Supplementary Table 1: Primary antibodies used in the study and their dilutions.**

| Primary antibodies | Tested samples | Dilutions | Vendors | Cat No. |
|---|---|---|---|---|
| Goat anti-CGRP | Mouse TMJs and knees | 1: 1000 | Biorad | 1720-9007 |
| Mouse anti-CGRP | Rat knees and spines | 1: 200 | Abcam | ab81887 |
| Mouse anti-NF200 | Pig TMJs | 1: 400 | Sigma | N0142 |
| Rabbit anti-CD31 | Mouse TMJs and knees, rat TMJs | 1: 50 | ThermoFisher | PA5-16301 |

CGRP: calcitonin-gene related peptide

NF200: neurofilament 200

CD31: cluster of differentiation 31, or platelet endothelial cell adhesion molecule (PECAM-1)

TMJs: temporomandibular joints.

**Supplementary Table 2: Timeline of joint processing, immunostaining, and clearing.**

| Joints / Protocols | Mouse | | Rat | | | Pig |
|---|---|---|---|---|---|---|
| | TMJs | Knees | TMJs | Knees | Spines | TMJs* |
| Decalcification | 3 | 3 | 7 | 7-10 | 7-10 | N/A |
| Permeabilization | 7 | 7 | 7 | 7 | 7 | 7 |
| Delipidation | 1-2 | 1-2 | 1-3 | 1-3 | 1-3 | 2-3 |
| Decolorization | 1 | 1 | 1 | 1 | 1 | 1 |
| GAGs extraction | 3 | 3 | 3 | 3 | 3 | 3 |
| Immunostaining | 15 | 15 | 23 | 23 | 23 | 23 |
| Clearing | 1 | 1 | 2-3 | 2-3 | 2-3 | 2-3 |
| Total time (days) | 31-32 | 31-32 | 44-47 | 44-50 | 44-50 | 38-42 |

GAGs: glycosaminoglycans.

TMJs: temporomandibular joints.

N/A: not applicable.

*Pig TMJ samples contain only the TMJ disc and surrounding soft tissues, such as retrodiscal tissues, so the decalcification step was not applied.

### Supplementary Notes:

#### Supplementary Note 1: Antibody verification and compatibility validation

Antibodies used in this study were listed in Supplementary Table 1. We first verified their specificity through 2D immunostaining. Mouse and rat spinal cords and skins and pig optical nerves were used to screen anti-calcitonin gene-related peptide (CGRP), CD31, and neurofilament 200 antibodies. Fresh samples were harvested from animals at the conclusion of other IACUC-approved research projects, fixed with 10% formalin at 4ºC for two days, and then washed in 1x PBS. Then, fixed samples were incubated in sucrose solutions, snap froze and sectioned into 10-15 μm frozen sections with a cryostat (CM1510S, Leica Microsystem, Inc., Exton, PA). Tissue sections were then washed in PBS to remove the O.C.T. (optimum cutting temperature) compound, permeabilized with 0.1% Triton X-100 in PBS, blocked with 10% donkey serum for 1 hour, and immunostained with primary antibody solution at 4ºC overnight. Next, the slides were washed and incubated in a secondary antibody solution at room temperature for 2 hours. Slides were washed, stained with a DAPI solution (R37606, ThermoFisher, Waltham, MA), and mounted with Fluoroshield (F6182, Sigma, St. Louis, MO) and a cover glass. Negative samples were defined as the sections only stained with the secondary antibody solution. Slides were imaged with a Leica TCS-SP5 confocal microscope (Leica Microsystem, Inc., Exton, PA) with 20x and 40x objectives. DAPI signal was captured with a 405 nm laser and emission filter range of 380-450 nm; signals from Alexa Fluor 488 dye were imaged with a 488 nm laser and emission filter of 490-540 nm; signals from Alexa Fluor 647 dye were detected with a 633 nm laser and emission filter range of 665-695 nm.

Our musculoskeletal joint immunostaining and clearing technique (MUSIC) method includes methanol pretreatment for delipidation and bleaching and uses methanol as the dehydration agent. An antibody validation experiment was performed as described in the literature to ensure the compatibility of the antibodies used in this study with methanol. Frozen sections were incubated in 100% methanol at room temperature for three hours and then rehydrated and proceeded with the immunostaining procedure described above.

#### Supplementary Note 2: Whole joint immunostaining validation

Our MUSIC method used multiple chemical agents to facilitate the deep permeabilization of the joint samples and to achieve whole joint mapping of neurovascular structures. To evaluate the performance of our method on deep immunostaining, we first immunostained the whole joint using our protocol with a secondary antibody with Alexa Fluor 647 dye (Supplementary Fig. 2a). Then, instead of processing the joints for tissue clearing, we snap-froze the joint and cut the joint into 10-15 μm sections for 2D immunostaining. We stained the sections with the same primary antibody used in the whole joint staining but used a different secondary antibody with Alexa Fluor 488 dye (Supplementary Fig. 2a). Slides were finally imaged with a confocal microscope by acquiring fluorescence signals from both channels of 488 and 647 dyes. As the antibody can directly and efficiently label tissue sections, the 2D immunostaining provides a ground truth to evaluate the efficiency of whole joint immunostaining. One expects to see overlap of the fluorescence signals from the whole joint staining (647 dye channel) and 2D section staining (488 dye channel) for the desired whole joint staining.

**Supplementary Movies:**

**Supplementary Movie 1: 3D neurovascular mapping in mouse TMJ.** The TMJ condyle, disc, and fossa were segmented and rendered in grey, translucent light blue, and translucent light brown, respectively.

**Supplementary Movie 2: 3D neurovascular mapping in wild-type mouse TMJ.** The TMJ condyles were segmented and rendered in grey for spatial reference.

**Supplementary Movie 3: 3D neurovascular mapping in *Prg4*$^{-/-}$ mouse TMJ.** The TMJ condyles were segmented and rendered in grey for spatial reference.

**Supplementary Movie 4: 3D vascular structures within a wild-type mouse TMJ condyle head.** The TMJ condyle was segmented and rendered in translucent color for spatial reference.

**Supplementary Movie 5: 3D vascular structures within a *Prg4*$^{-/-}$ mouse TMJ condyle head.** The TMJ condyle was segmented and rendered in translucent color for spatial reference.

**Supplementary Movie 6: 3D neurovascular mapping in Sham mouse TMJ.** The TMJ condyles were segmented and rendered in grey for spatial reference.

**Supplementary Movie 7: 3D neurovascular mapping in FMO_12d mouse TMJ.** The TMJ condyles were segmented and rendered in grey for spatial reference.

**Supplementary Movie 8: 3D neurovascular mapping in FMO_26d mouse TMJ.** The TMJ condyles were segmented and rendered in grey for spatial reference.

**Supplementary Movie 9: 3D neurovascular mapping in FMO_58d mouse TMJ.** The TMJ condyles were segmented and rendered in grey for spatial reference.

**Supplementary Movie 10: 3D neurovascular structures in wild-type mouse knee joint.** The femur and tibia bones were segmented and rendered in dark grey for spatial reference.

**Supplementary Movie 11: 3D neurovascular structures in *Prg4*$^{-/-}$ mouse knee joint.** The femur and tibia bones were segmented and rendered in dark grey for spatial reference.

**Supplementary Movie 12: 3D vascular structures in rat TMJ.**

**Supplementary Movie 13: 3D neural structures in rat knee joint.**

**Supplementary Movie 14: 3D neural structures in rat spine.**

**Supplementary Movie 15: 3D neural structures in pig TMJ disc.**
